# Relationship between melanoma vemurafenib tolerance thresholds and metabolic pathway choice and Wnt signaling involvement

**DOI:** 10.1101/2025.03.06.641924

**Authors:** Pratima Nangia-Makker, Madison Ahrens, Neeraja Purandare, Siddhesh Aras, Jing Li, Katherine Gurdziel, Hyejeong Jang, Seongho Kim, Malathy P Shekhar

## Abstract

Vemurafenib constitutes an important therapeutic for *BRAFV600* mutant melanomas, but despite high initial response rates, resistance to BRAF and MEK inhibitors quickly develops. Here, we performed an integrative analysis of metabolomic consequences and transcriptome alterations to uncover mechanisms involved in adaptive vemurafenib resistance (VemR) development and their relationship with vemurafenib tolerance thresholds. We developed *BRAFV600E* isogenic models of VemR utilizing M14 and A2058 lines, and patient-derived melanomas with V600E or normal BRAF to verify vemurafenib selectivity. MEK or PI3K inhibitors only partially inhibited VemR cell proliferation, indicating cross-resistance to these inhibitors. MITF and β-catenin levels were induced and treatment with Wnt/β-catenin inhibitor ICG-001 restored vemurafenib sensitivity with concomitant reductions in β-catenin-regulated gene expressions, phospho-ERK1/2, and VemR-induced mitochondrial mass and respiration. Targeted metabolite, MitoPlate-S1, Mito-stress and transcriptome/metabolomic analysis showed that melanoma cells with elevated vemurafenib tolerance thresholds such as A2058 VemR cells utilize Wnt/β-catenin signaling for mitochondrial metabolism while VemR cells with low tolerance such as M14 VemR cells rely on Wnt/β-catenin signaling for pentose phosphate pathway. Pathways associated with cytokine-cytokine receptor, ECM receptor, and neuroactive ligand receptor interactions were similarly enriched in *BRAFV600E* patient-derived melanoma as M14 and A2058 cells whereas distinct pathways involving cell cycle, DNA replication, Fanconi anemia and DNA repair pathways are upregulated in wild type BRAF expressing patient derived melanoma. These data show for the first time that the metabolic pathway choices made by VemR BRAF mutant melanomas are controlled by vemurafenib tolerance and endurance thresholds and Wnt/β-catenin signaling plays a central role in coordinating expression of genes controlling VemR and metabolic pathway shifts.

## Introduction

Malignant melanoma is a deadly form of skin cancer with exposure to ultraviolet radiation being a major risk factor for melanoma development. The 5-year survival rate for patients with metastatic melanoma is less than 25%.^1^ BRAF, a serine-threonine protein kinase, acting in the RAS-RAF-MEK-ERK signaling pathway, is mutated in approximately 65% of malignant cutaneous melanomas, and >90% of the cases involve V600 mutations.^2^ Approximately 15-20% of melanomas have mutations in *NRAS* while ∼6% of melanomas have normal BRAF.^3^ Vemurafenib and dabrafenib are FDA-approved BRAF inhibitors (BRAFi) for treating non-resectable melanomas with *V600E/K BRAF* mutation. Despite high initial response rates, acquired resistance to these drugs is common, and most patients relapse within months. The advent of immune checkpoint inhibitors (ICIs) have produced longer-lasting responses; however, > 75% of patients experience limited therapeutic benefit and initial ICI-responders develop disease progression. ^4,5^ Emerging evidence supports combining RAS–RAF-ERK pathway inhibitors such as vemurafenib with ICIs for improving therapeutic response, indicating BRAFi(s) continue to constitute an important armamentarium for metastatic melanoma treatment. Resistance to BRAFi(s) is linked to nonmutational adaptation of melanoma cells to the drug which is initiated during the early phase of treatment and continues through the tolerant phase of drug treatment.^6^ This provides ample opportunities for triggering or resetting pathways that augment BRAFi insensitivity such as reactivation of MAPK/ERK1/2 pathway, activation of PI3K/Akt signaling, and alterations in cellular metabolism.

Melanoma malignancy is associated with elevated glycolytic activity and lower mitochondrial respiration even under normoxic conditions,^7^ which is referred as the Warburg effect, a hallmark of many cancer types.^8^ *BRAFV600E* mutation is associated with metabolic reprogramming via induction of ERK1/2 hyperactivation and promotion of glycolytic enzymes enrichment.^9,10^ Compared to normal cells, cancer cells exhibit increased dependency on aerobic glycolysis, and fatty acid and nucleotide synthesis to sustain cell proliferation. ^11^ As an adaptive response to combat drug induced metabolic stress, BRAFi(s) reverse the Warburg effect, thus reducing glycolytic activity and stimulating mitochondrial biogenesis through the MITF-PGC1α pathway. ^12,13^ Accumulating evidence links altered metabolism with Wnt signaling, and aberrant activation of both the canonical (β-catenin dependent) and noncanonical (β-catenin independent) mechanisms of Wnt signaling contribute to neoplastic proliferation, metastatic dissemination, and therapy resistance.^14,15^ Wnt ligands WNT1, WNT2, WNT3, WNT3A, WNT8A, WNT8B, WNT10A and WNT10B activate the canonical pathway while WNT4, WNT5A, WNT5B, WNT6, WNT7A, WNT7B and WNT11 activate non-canonical Wnt-signaling,^16,17^ and the two pathways network and crosstalk to regulate one another. ^18,19^ The predominant reasons for upregulation of Wnt/β-catenin signaling is overexpression of canonical Wnt ligands or downregulation of Wnt antagonists (DKKs, WIF1, sFRP1/2) as β-catenin stabilizing mutations are rare in melanomas. Canonical Wnt/β-catenin signaling plays an important role in transcriptional activation of MITF, a lineage-specific transcription factor critical for melanocyte differentiation,^20^ and Wnt/β-catenin, MAPK/ERK and PI3K/AKT signaling pathways converge to regulate nuclear transport and transcriptional activity of MITF. ^21^ This suggests that canonical Wnt signaling may play a direct role in metabolic mitochondrial reprogramming via regulation of MITF mediated effects on mitochondrial biogenesis and respiration in BRAF mutant melanomas.

To establish the role of Wnt/β-catenin/MITF in metabolic reprogramming induced by adaptive vemurafenib resistance (VemR), we established BRAF mutant isogenic models of human metastatic melanoma with variant levels of VemR and performed an integrated analysis of the metabolic consequences and transcriptome alterations induced by adaptive VemR. Our data showed that acquisition of vemurafenib tolerance is associated with upregulation of Wnt/β-catenin signaling regardless of drug tolerance thresholds. Inhibition of β-catenin activity with the small molecule antagonist ICG-001 decreases VemR-induced mitochondrial mass and basal and maximal oxygen consumption rates. Our results from targeted metabolome and MitoPlate S-1 substrate utilization analysis revealed a critical role for β-catenin signaling in activating TCA cycle, a central hub in mitochondrial OXPHOS, for meeting the energy needs of A2058 VemR cells as compared to their sensitive parental counterparts. Our data also showed that unlike A2058 Verme cells, M14 VemR cells that have a lower threshold for vemurafenib tolerance utilize the pentose phosphate pathway which is also sensitive to β-catenin inhibition. Analysis of metabolites in BRAF wild type (Mel-14-089) and mutant (Mel-14-108) patient-derived melanoma cells confirmed the selectivity of vemurafenib as these metabolic pathways were perturbed only in BRAF mutant Mel-14-108 cells. Pathway analysis of transcripts showed similar enrichment of pathways associated with cytokine-cytokine receptor interaction, ECM receptor interaction, and neuroactive ligand receptor interaction in BRAF mutant Mel-14-108 PDX as in M14 and A2058 models, whereas pathways associated with cell cycle, DNA replication, Fanconi anemia and DNA repair pathways were enriched in wild type BRAF Mel-14-089 patient-derived melanoma, revealing distinct regulations by mutant and wild type BRAF. These data provide strong support for metabolic pathway choices triggered by vemurafenib sensitivity levels and the general therapeutic potential of targeting β-catenin oncogenic activity.

## Materials and Methods

### Cell Lines and cell culture

Human metastatic melanoma cells A2058 and M14 were obtained from American Type Culture Collection (ATCC, Manassas, VA) and the National Cancer Institute (Bethesda, MD), respectively. A2058 and M14 cells were cultured in Dulbecco’s modified Eagle’s medium (DMEM)/F12 medium supplemented with 5% fetal bovine serum at 37 °C, 5% CO_2_.^22^ The authenticated cell lines were used within 5–10 passages. Isogenic cells resistant to vemurafenib were generated by using a pulsed treatment strategy. M14 or A2058 cells were exposed to vemurafenib at their respective IC50 starting doses of 10 or 250 nM. Cells were treated for several days and then allowed to recover in drug free medium. Doses were gradually increased with similar pulsed on-off rounds. Cell survival was assessed every 3-5 weeks to confirm sustained and increased tolerance thresholds for vemurafenib compared to their parental counterparts, and IC50 values were determined regularly. Vemurafenib resistant (VemR) M14 or A2058 cells were maintained in DMEM/F12 medium supplemented with 5% FBS containing 100 nM or 5 μM vemurafenib, respectively. Patient-derived brain metastatic melanoma cells Mel-14-089 and Mel-14-108 were grown in DMEM/F12 media supplemented with 10% fetal bovine serum, non-essential amino acids, and gentamicin (Millipore Sigma, St. Louis, MO, USA) at 37 °C, 5% CO_2_. The details of Mel-14-089 and Mel-14-108 isolation from metastatic brain tumors were previously described.^23^ Whereas Mel-14–108 cells express *BRAF V600E*, Mel-14–089 cells express wild type BRAF. Acquisition and use of patient derived samples were approved by the Wayne State University Institutional Review Board and written informed consent was obtained from each patient prior to enrollment (IRB 111610MP2E; Protocol # 1011009008).

### Cell Survival and Colony Forming Analyses

Metastatic melanoma cells M14, A2058, their VemR counterparts M14 VemR and A2058 VemR, and patient derived Mel-14-089 and Mel-14-108 cells were seeded at 7,500 cells/well in 96-well plates in quadruplicates and treated with 0-10 µM vemurafenib or ICG-001 (Selleck Chemicals, Houston, TX, USA). Cell viability was assessed at 72 h post treatment by MTT assays. Experiments were performed in triplicates and results presented are representative of three independent experiments. For evaluation of colony forming potentials, parental M14 and A2058 cells were treated overnight with 0-100 nM or 0-500 nM vemurafenib, respectively, and M14 VemR and A2058 VemR cells were treated with 100 nM or 5 µM vemurafenib, respectively. Cells were trypsinized and reseeded in quadruplicates at 250 cells/well in 24-well plates in drug-free media. Colonies were allowed to form for two-three weeks and detected by crystal violet staining. Colonies were quantified using GelCount™ Oxford Optronix and CHARM algorithm with a minimum diameter of 100 μm set as the threshold for colony classification. Colony forming efficiencies were expressed relative to the untreated control cells.

### Western Blot Analysis

Whole cell lysates were prepared as previously described ^24^ from parental and VemR M14 and A2058 cells, and patient derived Mel-14-089 or Mel-14-108 cells treated with vemurafenib or vehicle. 50-100 µg of protein were subjected to 4-20% gradient SDS-PAGE and western blot analysis of phospho-ERK1/2 (Cell Signaling, Danvers, MA), ERK1/2 (Cell Signaling), Ser473phospho-AKT (Cell Signaling), AKT (Cell Signaling), β-catenin (Santa Cruz Biotechnology, Dallas, TX), Melan A (Dako Corp., Santa Clara, CA), Vimentin (Dako Corp.), MITF (Abcam, Cambridge, MA), Total OXPHOS human antibody cocktail (Abcam, Waltham, MA), Tomm 20 (Santa Cruz Biotechnology), ABCB5 (Abcam), β-actin and α-tubulin (Sigma-Aldrich Chemicals, St. Louis, MO). Protein levels relative to the loading control α-tubulin or β-actin were quantified by ImageJ version 1.53 (NIH, Bethesda, MD).

### Chemotaxis analysis

A2058 VemR cells were treated overnight with 5 µM vemurafenib alone or in combination with 0-10 µM ICG-001, and subjected to Boyden chamber chemotaxis assays as previously described.^25^ Cells that migrated across the membrane were fixed and stained using Diff-Quik stain set (Baxter, Deerfield, IL). The staining intensities were quantified using ImageJ version 1.53.

### Dual-Luciferase Reporter Assay

To evaluate the transcriptional activity of endogenous TCF/β-catenin, parental and VemR M14 and A2058 cells were transiently cotransfected with 1.0 µg of pTOP/FLASH or pFOP/FLASH (Upstate Biotech, Lake Placid, NY) and 100 ng of Renilla luciferase pRL-TK (Promega, Madison, WI) using Metafectene (Biontex Laboratories GmbH, Munich, Germany). Luciferase activities were measured as previously described using the Promega Dual Luciferase Assay system.^26^ Absolute promoter firefly luciferase activity was normalized against Renilla luciferase activity to correct for transfection efficiency. Triplicate dishes were assayed for each transfection and at least three transfection assays were performed.

### Mitochondria imaging analysis

Parental and VemR M14 or A2058 cells cultured on coverslips were treated overnight with vemurafenib (M14, 10 nM; M14 VemR, 100 nM; A2058, 250 nM; A2058 VemR, 5 µM) alone or in combination with 5 µM ICG-001. Cells were washed with serum free media before incubation with 50 nM MitoTracker DeepRed FM (Molecular Probes, Eugene, OR, USA), a mitochondrial membrane potential-dependent dye, at 37°C for 30 min. Cells were washed, fixed, and mounted in Slowfade containing 4′,6-diamidino-2-phenylindole (DAPI) to counterstain nuclei. Images were captured on Olympus BX40 microscope equipped with a Sony high resolution/sensitivity CCD video camera and CellSens software. The percentage of cells with active mitochondria and the integrated staining densities reflecting active mitochondrial mass were determined using ImageJ tool and scored from at least 30-50 cells/field and three to five fields.

### Analysis of sugar utilization flexibility

To compare sugar utilization flexibility and survival of parental and VemR M14 or A2058 cells, cells were cultured in media containing high or low glucose or galactose. Dialyzed fetal bovine serum (5%) was used for all experiments. The compositions of the media were: high glucose medium: DMEM deprived of glucose (Invitrogen A14430) supplemented with 25 mM D-glucose, 0.5 mM sodium pyruvate, 2.5 mM L-glutamine, 5% FBS, and penicillin-streptomycin (500 μg/ml final concentration). Low glucose medium: DMEM deprived of glucose supplemented with 5 mM D-glucose, 0.5 mM sodium pyruvate, 2.5 mM L-glutamine, 5% FBS, and penicillin-streptomycin. Galactose medium: DMEM deprived of glucose supplemented with 25 mM D-galactose, 0.5 mM sodium pyruvate, 2.5 mM L-glutamine, 5% FBS, and penicillin-streptomycin. Prior to the experiment, cells cultured in high glucose DMEM/F12 media were washed with PBS followed by incubation for 8-10 hours in sugar-free medium to facilitate glucose depletion. Cells were trypsinized, pelleted, and rinsed thrice with PBS. After washing, cell pellets were resuspended in high glucose, low glucose or galactose media, and seeded at 2×10^4^ cells/well in quadruplicates in 96-well plates. Cell viability was measured at 72 to 96 h by MTT assay, and the results were expressed as mean ± S.D. relative to corresponding high glucose condition. At least three independent assays were performed. To determine the functional involvement of β-catenin on sugar utilization, M14 and A2058 VemR cells were seeded under high glucose and galactose conditions in 96-well plates and treated with 1-10 µM ICG-001. Cell viability was measured at 72 to 96 h by MTT assay as described above.

### Cellular bioenergetics analysis

Oxygen consumption rates (OCR) of parental and VemR M14 and A2058 cells cultured in glucose or galactose containing media or treated with ICG-001 were measured using the XFe24 flux analyzer and the Cell Mito Stress Test kit (Agilent Technologies) according to the manufacturer’s instructions. Cells were seeded at a density of 3 × 10^4^ cells/well in quintuplicates in Seahorse XFe24 24-well plates at 37°C. Global mitochondrial parameters, i.e., basal respiration, ATP production, maximal respiratory capacity, and non-mitochondrial respiration were determined by sequential injection of ATPase inhibitor oligomycin (1 μM), an uncoupler of the respiratory chain carbonyl cyanide-4-(trifluoromethoxy)phenylhydrazone (FCCP, 1 μM), and ETC complex I and III inhibitors, rotenone (0.5 μM) and antimycin A (0.5 μM), respectively. Data were analyzed using the XF software (Agilent Technologies).

### Immunofluorescence staining

M14 VemR and A2058 VemR cells were seeded on coverslips and treated overnight with 100 nM or 4 µM vemurafenib, respectively, or in combination with 5 µM ICG-001. Cells were fixed with 10% phosphate buffered formalin, permeabilized with methanol:acetone (1:1, v/v), and immunostained with β-catenin antibody (SantaCruz Biotechnology) and corresponding Texas Red-conjugated secondary antibody (Molecular Probes, Eugene, OR). Nuclei were counterstained with DAPI. Images were collected on an Olympus BX60 microscope equipped with a Sony high resolution/sensitivity CCD video camera and processed using CellSens software.

### Targeted metabolomics

Steady-state levels of metabolites in parental and VemR M14 and A2058 cells were quantitatively profiled using an established LC-MS/MS-based targeted metabolomics platform, which measures 254 metabolites involved in major human metabolic pathways.^27^ All LC-MS/MS analyses were performed on an AB SCIEX QTRAP 6500 LC-MS/MS system, which consists of a SHIMADZU Nexera ultra-high-performance liquid chromatography coupled with a triple quadrupole/linear ion trap mass spectrometer. Analyst 1.6 software was used for system control and data acquisition, and MultiQuant 3.0 software was used for data processing and quantitation. Metabolites in parental and VemR M14 and A2058 cells were extracted with ice-cold 80% methanol and subjected to LC-MS/MS analyses as previously described.^27^ Metabolite concentrations were normalized to cellular protein. Statistical analysis and pathway analysis were conducted using MetaboAnalyst (www.MetaboAnalyst.ca, version 6.0).^28^ Comparisons of individual metabolites between the groups were performed using unpaired t-tests with FDR correction applied using the Benjamini-Hochberg method.

### Mitochondrial and cytosol metabolism analysis

Alterations in metabolism inside the mitochondria and/or cytosol induced by acquired vemurafenib resistance were evaluated using MitoPlate S-1 assay (Biolog Inc., Hayward, CA). MitoPlate S-1 plate contains 3 sets of precoated 31 substrates of mitochondrial or glycolytic metabolism plus controls, and allows for interrogation and characterization of substrate metabolism rates. The assay was performed according to the manufacturer’s directions. Briefly, the precoated wells were incubated for 1 h at 37°C with 30 µl of Biolog mitochondrial assay solution (MAS) containing redox dye MC (Biolog Inc) and 100 µg/ml saponin. Parental and VemR M14 or A2058 cells were resuspended in Biolog MAS at 1×10^6^ cells/mL and 30 µl of cell suspensions were added to the preincubated wells. In some cases, cells were treated with 1 µM ICG-001 to determine the effect of β-catenin activity inhibition on mitochondrial and cytoplasmic metabolism. MitoPlates were immediately loaded into BioTek Synergy2 microplate reader to perform kinetic measurements at 590 nm at 10 min intervals for 4 h at 37°C. Data were normalized using the "No substrate" well (A1) required for the rows A to H (cytoplasmic substrates rows A and B; mitochondrial substrates rows C-H). Data were analyzed using R which creates a scatterplot of the initial rate values between conditions of interest.

### Whole Genome Expression Analysis by RNA-seq

To identify transcripts that are affected by VemR acquisition, parental M14 and A2058 cells and their VemR resistant counterparts were subjected to bulk RNA-seq analysis. To determine the impact of BRAF mutation on the transcriptome, metastatic melanoma PDXs with BRAF mutation (Mel-14-108) or wild type BRAF (Mel-14-089) were treated with 10 nM or 4 µM vemurafenib, respectively, and similarly subjected to bulk RNA-seq analysis. Total RNA was isolated using the Trizol reagent kit (Invitrogen, Carlsbad, CA), and RNA-seq libraries were prepared using Lexogen’s QuantSeq 3’mRNA-seq Library Prep Kit (FWD for Illumina) from 200 ng of DNase I treated RNA. The barcoded libraries were multiplexed (in batches of 27) at equimolar concentrations and sequenced with 50 bp reads on an Illumina NovaSeq 6000. RNA-seq analysis was conducted through the Genome Sciences Core at Wayne State University. After data were demultiplexed using Illumina’s CASAVA 1.8.2 software, reads were aligned to the human reference genome (Build hg38) ^29^ and tabulated before analysis with R/Bioconductor package edgeR (version 3.36.0).^30^ For differential gene expression analysis, the edgeR function ‘glmQLFTest’ was used. FDR was computed using Benjamini-Hochberg method, ^31^ and heatmap and hierarchical clustering were carried out using the R function Heatmap in the R package ComplexHeatmap (version 2.22.). Differentially expressed genes (DEGs) between parental and VemR counterparts were detected using 5% FDR and fold-change (FC) of ≥1.5. iPathwayGuide Analysis (Advaita Bioinformatics) was performed to obtain biological information on the pathways perturbed by vemurafenib resistance acquisition.

### Real-Time Reverse transcriptase (RT)-PCR Analysis

The accuracy of sequencing data was validated by quantitative RT-PCR analysis. qRT-PCR analysis of β-catenin, MITF-M, TYRP1, ABCB5, RBKS and GAPDH reference control expressions in M14 VemR and A2058 VemR cells were performed using DNase I treated total RNAs and Maxima SYBR Green/ROX qPCR mix (ThermoFisher, Waltham, MA). Primers for real-time RT-PCR analysis were as follows: β-catenin (accession number NM_001098209), 5’- ATACCACCCACTTGGCAGAC-3’ (forward) and 5’-GGAAGGTCTCCTTGGGACTC-3’ (reverse); TYRP1 (accession number NM_000550), 5’-ATGGCAACACGCCACAATTTGAG-3’ (forward) and 5’-CCCGTTGCAAAATTCCAGTAAG-3’ (reverse); MITF-M (MITF transcript isoform 4; accession number NM_000248), 5’- ATGCTGGAAATGCTAGAATATAATCACTATCAG-3’ (forward) and 5’- AGCCATGGGGCTGTTGGGTGC-3’ (reverse); RBKS (accession number BC017425.1), 5’- GCCGAGCCAAAGTGATGATATG-3’ (forward) and 5’-TGGAGAGGGTATAGAACTGGGG- 3’ (reverse); ABCB5 (accession number MK803370), 5’-GCTGAGGAATCCACCCAATCT-3 ’ (forward) and 5’-CACAAAAGGCCATTCAGGCT-3’ (reverse), and GAPDH (accession number NM_002046), 5’-AAATATGATGACACCAAGAAGG-3’ (forward) and 5’- TGAAGTCGGAGGAGACCAC-3’ (reverse). Amplification was performed in StepOne Plus Real-Time PCR system (Applied Biosystems), and the cycling conditions were 10 minutes at 95°C, and 35 cycles of 15 seconds at 95°C, 1 minute at 57°C, and 1 minute at 68°C.

### Statistical analysis

Comparisons between two groups were performed using unpaired t-tests, and for three or more groups, ANOVA was utilized along with Holm’s post-hoc analysis. Differential analyses for metabolomics and RNA-seq data were performed using unpaired t-tests or moderated t-tests, followed by FDR correction. The volcano plot was generated with FDR <0.05 and fold-change >1.5. The meta-analysis was performed based on Venn Diagrams and UpSet plots. Multi-omics analysis was conducted between metabolomics and RNA-seq data with significant pathways identified using metabolomics data. For each significant pathway, genes that were present in the pathway were extracted to construct a gene set. For each of the gene sets, heatmaps were generated followed by gene set enrichment analysis (GSEA). Experimental results are presented as the mean ± standard deviation (S.D.) or standard error of mean (S.E.M). Statistical significances were considered if P < 0.05. All statistical analyses were performed with GraphPad Prism, Microsoft Excel, and the statistical computing software R.

## Results

### Generation and characterization of isogenic models of vemurafenib resistant (VemR) melanoma cells

The melanoma lines, M14 and A2058, were chosen as they both carry *BRAF V600E* mutation. ^32^ To check their sensitivities to vemurafenib, a selective inhibitor of mutant BRAF, MTT assays were performed. The results show that the M14 cell line is more responsive to vemurafenib treatment compared to A2058 cells with IC50 values of 10 nM and 250 nM, respectively (Supplementary Fig. S1A and B). The results of MTT assays were further verified by colony forming assays. Colony forming potentials of M14 and A2058 cells were significantly inhibited with 5 nM (P<0.001) and 50 M (P<0.01) vemurafenib (Supplementary Fig. S1 C and D). The representative images of M14 and A2058 colonies captured by GelCount Oxford Optronix are shown in Supplementary Fig. 1E and 1F, respectively. Clinical trials using vemurafenib and other selective BRAFi have shown impressive results, however, despite initial therapeutic responses, development of acquired or adaptive resistance limits their clinical efficacy. To develop isogenic models of VemR, cells were exposed to gradually increasing concentrations of vemurafenib over several weeks. Cell survival was assessed every 3-5 weeks to verify increased tolerance thresholds for vemurafenib compared to their parental counterparts, and IC_50_ values were determined regularly. MTT assays confirmed the increased abilities of vemurafenib-adapted M14 and A2058 cells to tolerate vemurafenib with M14 VemR and A2058 VemR cells displaying IC50s of 0.1 µM and 5 µM, respectively, as compared to their parental counterparts with IC50s of 10 nM and 250 nM, respectively (Fig. 1A,B). M14 and A2058 VemR cells were continuously maintained in medium containing 100 nM or 5 µM vemurafenib, respectively.

**Figure 1.**
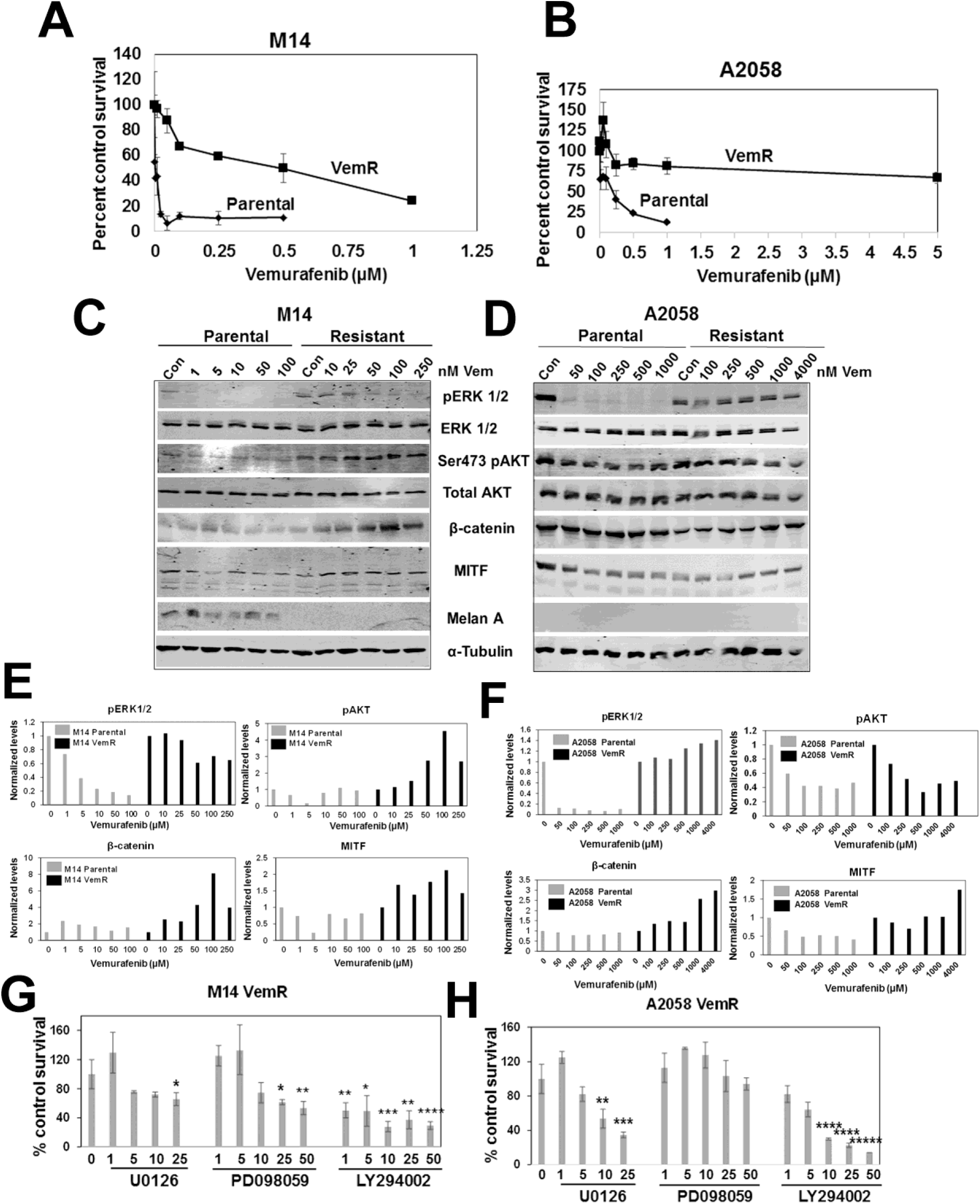
Characterization of isogenic melanoma models of vemurafenib resistance. (A,B) MTT assays of isogenic pairs of parental and vemurafenib resistant (VemR) M14 and A2058 cells. (C,D) Western blot analysis of the indicated proteins following treatment with the indicated doses of vemurafenib, and (E,F) quantification of the indicated protein levels from representative blots of M14 (E) and A2058 (F) isogenic pairs. (G,H) Cell survival analysis of M14 VemR (G) and A2058 VemR (H) treated with MEK (U0126, PD098059) and PI3K (LY294002) inhibitors by MTT assay. Results are expressed as mean ± S.D. (percent of control cell survival) from three independent experiments. *P<0.05; **P<0.01; ***P<0.001; ****P<0.0001; *****P<0.00001.

To establish whether the differences in vemurafenib sensitivities between the corresponding isogenic pairs correlated with differences in inhibition of ERK kinase, a direct downstream effector of BRAF, we evaluated the expression levels of phospho-ERK1/2. Whereas ERK phosphorylation was inhibited by vemurafenib treatment in parental M14 and A2058 lines (Fig. 1C,D), commensurate with their increased drug tolerability, phospho-ERK1/2 levels remained largely unaffected by vemurafenib in the resistant counterparts (Fig. 1C-F). Expression levels of Akt Ser473 phosphorylation were analyzed since PI3K/AKT pathway is implicated in VemR. Phospho-Ser473 Akt levels were upregulated or unaffected by VemR in M14 VemR or A2058 VemR cells, respectively, as compared to their parental controls (Fig. 1C-F).

Activation of Wnt/β-catenin signaling and MITF, a β-catenin transcriptional target, are implicated as drivers of BRAFi resistance in cancer cells carrying *BRAFV600E* mutation.^33,34^ To determine whether BRAF/Wnt/β-catenin crosstalk plays a role in adaptive VemR, we analyzed vemurafenib effects on β-catenin and MITF levels. While β-catenin levels were unaffected by vemurafenib in both M14 and A2058 parental lines (Fig. 1C-F), treatment with ≥50 nM or ≥ 1 µM vemurafenib induced 2-4-fold increases in β-catenin levels in M14 VemR and A2058 VemR cells, respectively (Fig. 1C-F). MITF protein levels were increased ∼2-fold by vemurafenib in VemR models, mirroring β-catenin profiles (Fig. 1C-F).

To further confirm the relevance of ERK and Akt activities in VemR, we analyzed the effects of MEK (U0126 and PD098059) and PI3K/Akt (LY294002) inhibitors on M14 VemR and A2058 VemR cell survival by MTT assays. Both VemR models were less sensitive to MEK inhibitors requiring 25 µM PD098059 and >10 µM U0126 to induce 50% inhibition of cell survival (Fig. 1G, H). Concentrations of >10 µM LY294002 were required to inhibit M14 VemR and A2058 VemR cell survival, suggesting a moderate role for PI3K/Akt activity in our models. Taken together, these data establish the validity of our VemR isogenic models and implicate a potential role for β-catenin signaling in VemR.

### β-catenin transcriptional activity inhibition sensitizes vemurafenib resistant melanoma cells

To determine if activation of β-catenin signaling contributes to increased tolerance to vemurafenib, we analyzed the effect of β-catenin transcriptional inhibitor, ICG-001, on vemurafenib sensitivity by MTT and colony forming assays. M14 VemR and A2058 VemR cells cultured in the presence of 100 nM or 5 µM vemurafenib, respectively, were treated with 0.1 – 5 µM ICG-001. Treatment with ICG-001 significantly decreased the survival of both M14 VemR (P<0.01) and A2058 VemR (P<0.05) cells, albeit more strongly in M14 VemR cells (Figure 2A, B). Phase-contrast microscopy confirmed ICG-001 induced cytotoxicity in VemR cells (Fig. 2C, D). Treatment with 5 or 10 µM ICG-001 abrogated M14 VemR and A2058 VemR colony formation, verifying the MTT assay results (Fig. 2E-H). Treatment with ML329, a small molecule inhibitor of MITF, significantly inhibited M14 VemR and A2058 VemR cell survival (P<0.01), supporting the role of MITF (a β-catenin transcriptional target) and β-catenin activity in VemR (Supplementary Fig. S2).

**Figure 2.**
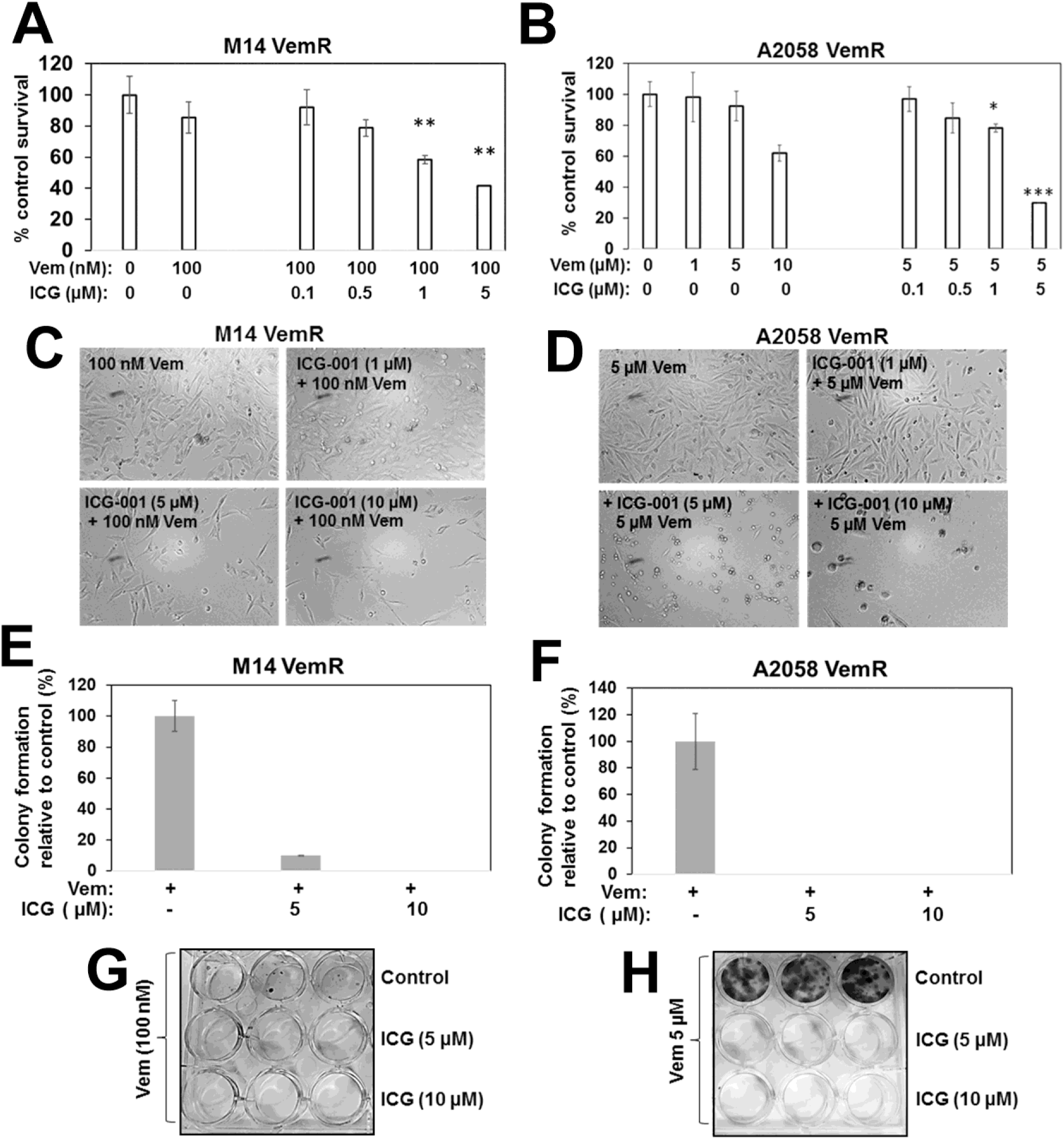
Sensitivity analysis of β-catenin inhibitor ICG-001 on vemurafenib sensitivities of M14 and A2058 VemR cells. (A,B) Evaluation of ICG-001 effects on survival of M14 VemR cells exposed to 100 nM vemurafenib (A) or A2058 VemR cells exposed to 5 µM vemurafenib (B) by MTT assays, and (C,D) corresponding phase-contrast micrographs. (E,F) ICG-001 effects on colony forming potentials of M14 VemR (E) and A2058 VemR (F) cells. Results are expressed as mean ± S.D. (percent of control colony formation efficiency) from three independent experiments. *P<0.05; **P<0.01; ***P<0.001. (G,H) Images of representative colonies captured using GelCount Oxford Optronix.

To determine whether ICG-001-induced sensitization of VemR cells resulted from combined inhibition of ERK and β-catenin signaling, we analyzed the effects of ICG-001 on phospho-ERK1/2, ERK1/2, β-catenin, MITF and vimentin (also a β-catenin transcriptional target) protein levels. Treatment with ICG-001 elicited noticeable decreases in phospho-ERK1/2, β-catenin, MITF and vimentin in both VemR models compared to cells treated with vemurafenib alone (Fig. 3A-D). ICG-001 treatment also reduced total ERK1/2 levels in A2058 VemR cells, suggesting potential impact of β-catenin inhibitor on ERK1/2 stability (Fig. 3A). TOP/Flash reporter assays showed ∼2-3-fold increases, respectively, in β-catenin dependent transcriptional activity in M14 VemR and A2058 VemR cells as compared to their respective parental counterparts (P<0.01), and treatment with ICG-001 significantly decreased TOP/Flash activity in M14 VemR (P<0.001) and A2058 VemR (P<0.01) cells (Fig. 3E). Consistent with TOP/Flash reporter data, real-time RT-PCR analysis showed ICG-001 significantly decreased expressions of β-catenin (P<0.05), MITF-M (P<0.001) and TYRP1 (P<0.001) in both VemR models (Fig. 3F, G). Boyden chamber assays showed that ICG-001 treatment decreased A2058 VemR cell migration in a dose-dependent manner as compared to cells treated with vemurafenib alone (Fig. 3H, I), corroborating β-catenin involvement in migration and invasion. Taken together, these data reveal a major role for Wnt/β-catenin signaling in persistent activation of ERK1/2 in VemR cell, and that inhibition of β-catenin transcriptional activity can disrupt this crosstalk and suppress BRAFi resistance.

**Figure 3.**
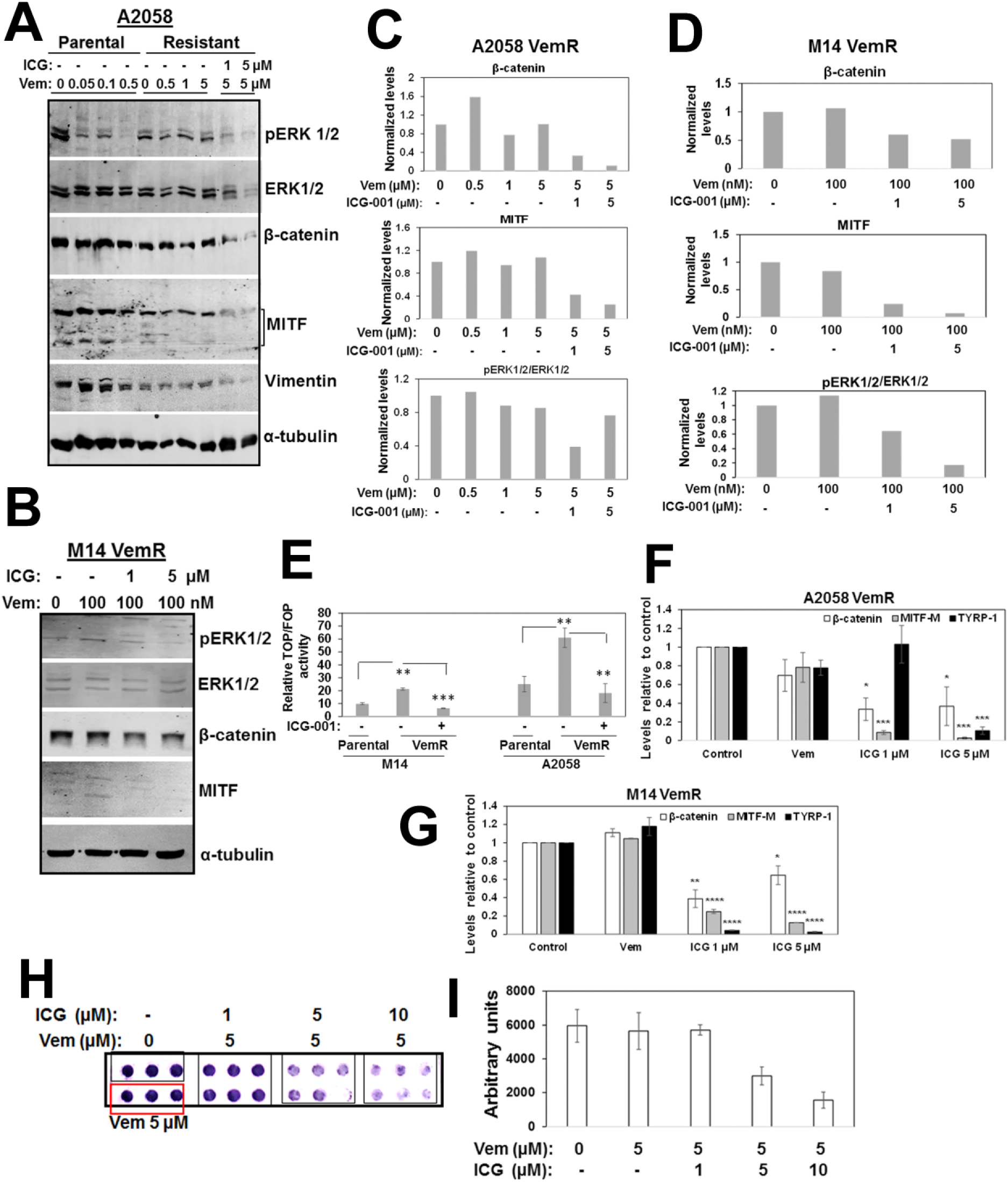
ICG-001 treatment downregulated β-catenin mediated transcriptional activity and β-catenin regulated genes. (A,B) Western blot analysis of the indicated proteins following treatment with vemurafenib or ICG-001 at the indicated concentrations, and (C,D) relative levels of the indicated proteins in A2058 VemR (C) and M14 VemR (D) cells. (E) Analysis of β-catenin transcriptional activity and ICG-001 regulation in parental and VemR M14 and A2058 isogenic cells. (F,G) Real-time quantitative RT-PCR analysis of β-catenin regulated gene expression in A2058 VemR (F) and M14 VemR (G) cells. (H) ICG-001 effects on A2058 VemR cell migration using Boyden Chamber, and (I) quantitation using Image J software. *P<0.05; **P<0.01; ***P<0.001; ****P<0.0001.

To determine the clinical relevance of BRAF/ERK and β-catenin signaling, we conducted studies using two patient derived brain metastatic melanomas, Mel-14-108 and Mel-14-089, that express *BRAFV600E* or wild type BRAF, respectively. Consistent with BRAF mutational status, MTT assays showed that Mel-14-108 cells are very sensitive to vemurafenib (IC50 ∼40 nM) whereas 50% of Mel-14-089 remain relatively unaffected even with 10 µM vemurafenib (Fig. 4A, B). These data are corroborated by western blot analysis that showed vemurafenib inhibited ERK1/2 phosphorylation in Mel-14-108 cells but not in Mel-14-089 cells (Fig. 4C). Concomitant with ERK1/2 inhibition, vemurafenib treatment also inhibited Akt Ser473 phosphorylation in Mel-14-108 cells while Akt remained insensitive to vemurafenib in Mel-14-089 cells. These data suggest a common link between ERK1/2 and Akt regulation and vemurafenib sensitivity in melanoma PDXs. To test whether β-catenin/BRAF crosstalk is affected by BRAF mutational status, we analyzed the effects of ICG-001 on Mel-14-108 or Mel-14-089 cell survival by MTT assays. Despite differences in vemurafenib responses, ICG-001 treatment inhibited proliferation of both melanoma PDXs (Fig. 4D, E). These data suggest a BRAF mutation-independent role for β-catenin in vemurafenib refractory melanomas and that melanomas with mutant or wild type BRAF could benefit from inhibition of β-catenin signaling.

**Figure 4.**
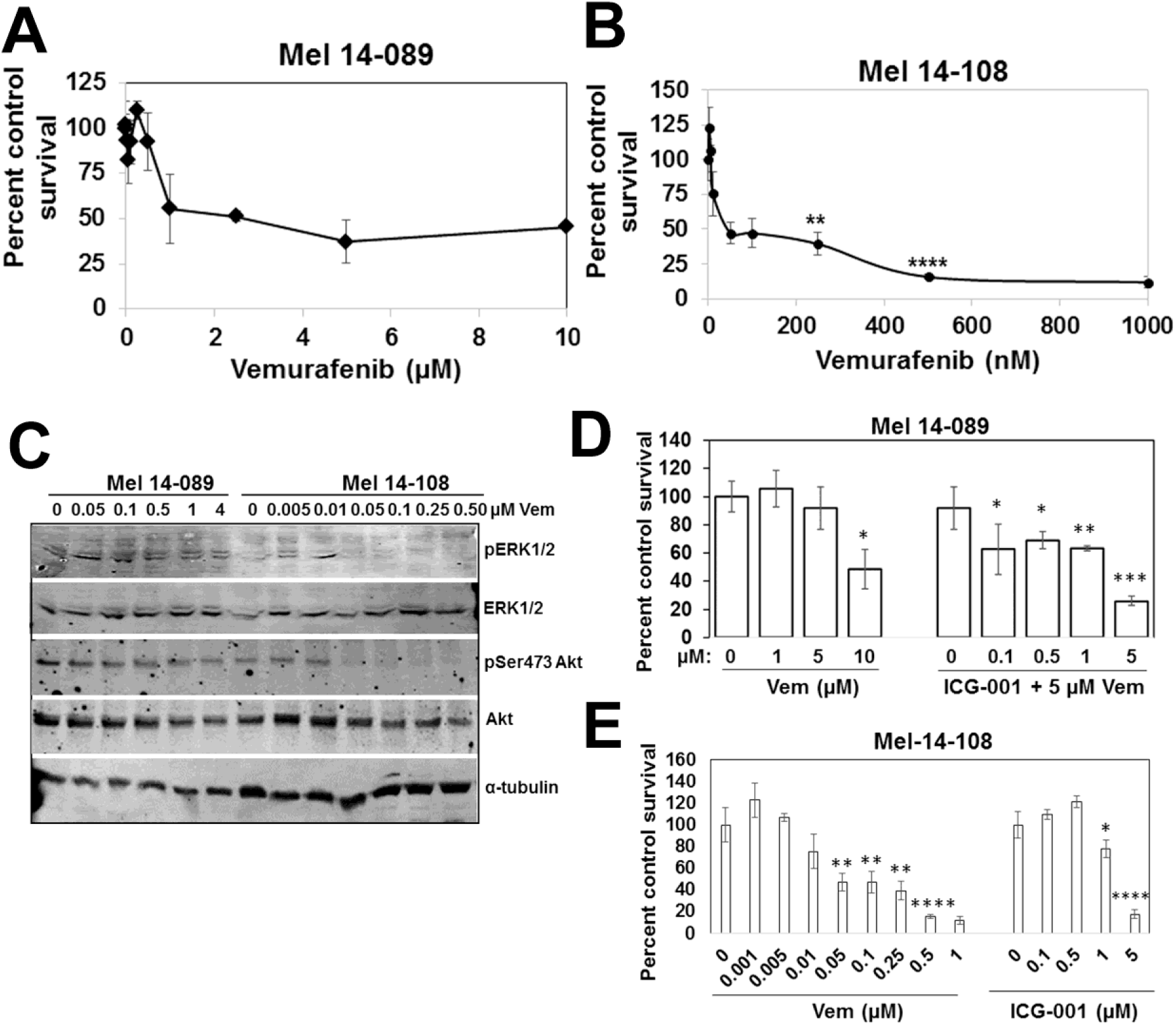
Vemurafenib sensitivity and ICG-001 effects on patient derived melanoma cells with wild type or mutant BRAF. (A,B) Evaluation of vemurafenib sensitivities of wild type BRAF Mel-14-089 (A) and V600E BRAF Mel-14-108 (B) cells by MTT assays. Results are expressed as mean ± S.D. relative to control cell survival from two independent experiments each performed in quadruplicates. (C) Western blot analysis of the indicated proteins in vemurafenib treated Mel-14-089 and Mel-14-108 cells. (D,E) ICG-001 treatment inhibits survival of both patient derived Mel-14-089 (D) and Mel-14-108 (E) melanoma cells. Results are expressed as mean ± S.D. relative to control cells from two independent experiments, each performed in quadruplicates. *P<0.05; **P<0.01; ***P<0.001, ****P<0.0001.

### VemR melanoma cells have elevated levels of mitochondrial mass and OXPHOS activity

WNT signaling functions in both glycolysis and mitochondrial respiration.^35^ Since our data implicated β-catenin and MITF in adaptive VemR, we next determined if adaptive VemR involves metabolic rewiring to mitochondrial metabolism. Mitochondrial mass and activity of parental and VemR cells were measured by staining with Mitotracker Deep Red FM, a mitochondrial potential dependent dye. Vemurafenib treatment of parental M14 and A2058 cells resulted in significant decreases in the percentage of cells with active mitochondria (P<0.05) and mitochondrial mass (P<0.01) as compared to untreated cells (Figure 5A-D). In contrast, M14 VemR and A2058 VemR cells displayed stronger Mitotracker Red staining intensities and mitochondrial mass that were unaffected by treatment with 100 nM or 5 µM vemurafenib, respectively (Figure 5A-D).

**Figure 5.**
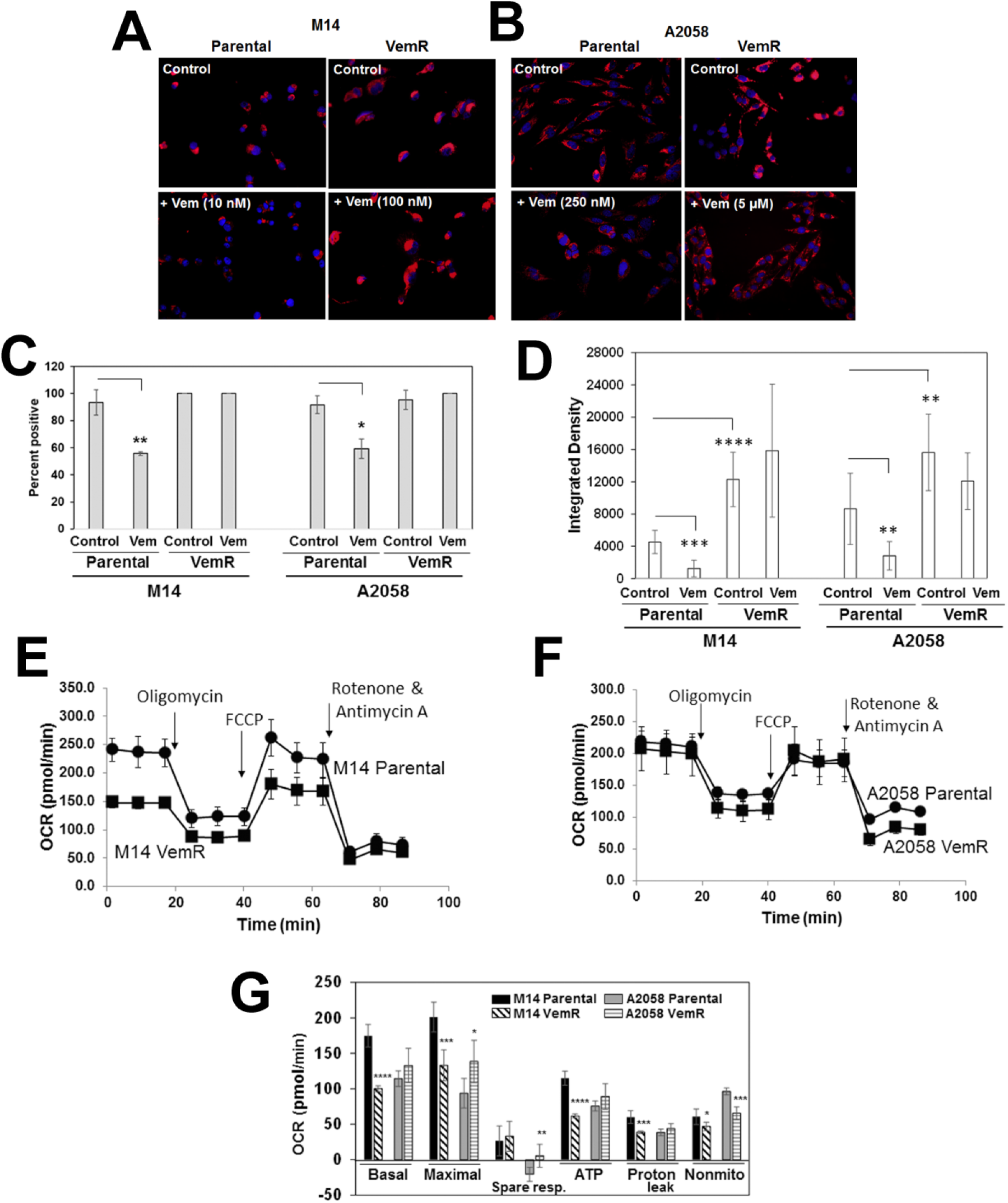
Vemurafenib resistant melanoma cells have increased mitochondrial mass and spare respiratory capacity. (A,B) MitoTracker DeepRed staining of parental and VemR M14 and A2058 cells. Original magnification ×100. (C) Percent of MitoTracker Red positive cells and (D) active mitochondrial area density determined using ImageJ. A minimum of 30-50 cells from three to five fields were scored and results expressed mean ± S.D. (E-G) Bioenergetics analysis of parental and VemR M14 (E) and A2058 (F) cells by Seahorse flux analyzer using mito stress kit. (G) Summary data calculated from the curves in E and F. *, P<0.05; **, P<0.01; ***, P<0. 001 ****; P<0.0001.

The oxygen consumption rate (OCR), an indicator of OXPHOS activity, was determined using the Seahorse XF Cell Mito Stress test under basal conditions and in response to sequential treatment with electron chain complex inhibitors (oligomycin, rotenone/antimycin A) and mitochondrial decouplers (FCCP). Mitochondrial functions were normalized to cell number and calculated based on the bioenergetics profiles that include basal respiratory capacity, maximal respiration, spare respiratory capacity and ATP production. The results in Fig. 5F and G show that parental and VemR A2058 cells have similar levels of basal OCR. However, maximal respiration, an indicator of reserve or spare respiratory capacity (after injection of the uncoupler FCCP), was significantly higher in A2058 VemR (P = 0.027) cells as compared to its parental counterpart. M14 VemR cells have significantly lower basal and maximal OCR compared to their parental counterparts (P<0.001) (Fig. 5E, G). However, although maximal respiration was lower in M14 VemR cells as compared to parental cells, the spare respiratory capacity of M14 VemR cells was slightly higher. although not significantly, suggesting its potential for coping with drug induced metabolic stress (Figure 5G). ATP-linked respiration was higher in A2058 VemR cells, reflecting its greater threshold of vemurafenib tolerance. These data are consistent with the greater Mitotracker Red staining intensities and mitochondrial mass in VemR cells and suggest that alterations in mitochondrial bioenergetics are commensurate with vemurafenib tolerance thresholds.

To gain insights into regulation of OXPHOS levels by vemurafenib in parental and VemR M14 and A2058 cells, we evaluated the levels of subunits of OXPHOS complexes II-V by western blot analysis. Among the four complexes, levels of UQCRC2 (ubiquinol-cytochrome c reductase), a core protein required for assembly of complex III, were marginally higher in both VemR models as compared to their parental counterparts. COX II (complex IV subunit) levels were also marginally induced in A2058 VemR, and no discernible differences in SDHB (complex II subunit) or ATP5A (complex V subunit) protein levels between the parental and VemR cells were observed (Supplementary Fig. S3).

### VemR melanoma cells have increased abilities to utilize galactose as the sole energy source

Our results thus far suggest that VemR acquisition involves gain in mitochondrial activity. To confirm these data, we tested the impact of glucose vs. galactose as the only oxidizable substrate on survival and bioenergetics. Parental and VemR M14 or A2058 isogenic cells were cultured in medium containing 25 mM (high) glucose, 5 mM (low) glucose or 25 mM galactose, and their survival under these conditions was assessed by MTT assays. Replacement of glucose with galactose forces cells to shift their energy source from cytosolic glycolysis to mitochondrial oxidative phosphorylation.^36^ Compared to parental M14 cells that were significantly growth inhibited in low glucose or galactose media (P<0.001), their VemR counterpart survived significantly better (∼5-fold) under these conditions (P<0.001), suggesting gain or improvement in mitochondrial function. A2058 VemR cells showed similar increases in ability to utilize galactose, albeit smaller magnitude (∼2-fold), as compared to their parental counterparts (P<0.05; Figure 6A). Since unlike M14 VemR cells, A2058 VemR cells performed only slightly better than their parental counterparts under low glucose or galactose conditions, we posit that the small differences in magnitude of metabolic shifts between parental and VemR A2058 cells potentially reflect their intrinsically greater thresholds for vemurafenib tolerance. Since parental A2058 cells are inherently more tolerant of vemurafenib (IC50 250 nM) compared to parental M14 cells (IC50 10 nM), we speculate that parental A2058 cells are already utilizing both glycolytic and mitochondrial pathways for their energy needs while M14 VemR cells may be forced to use mitochondria.

**Figure 6.**
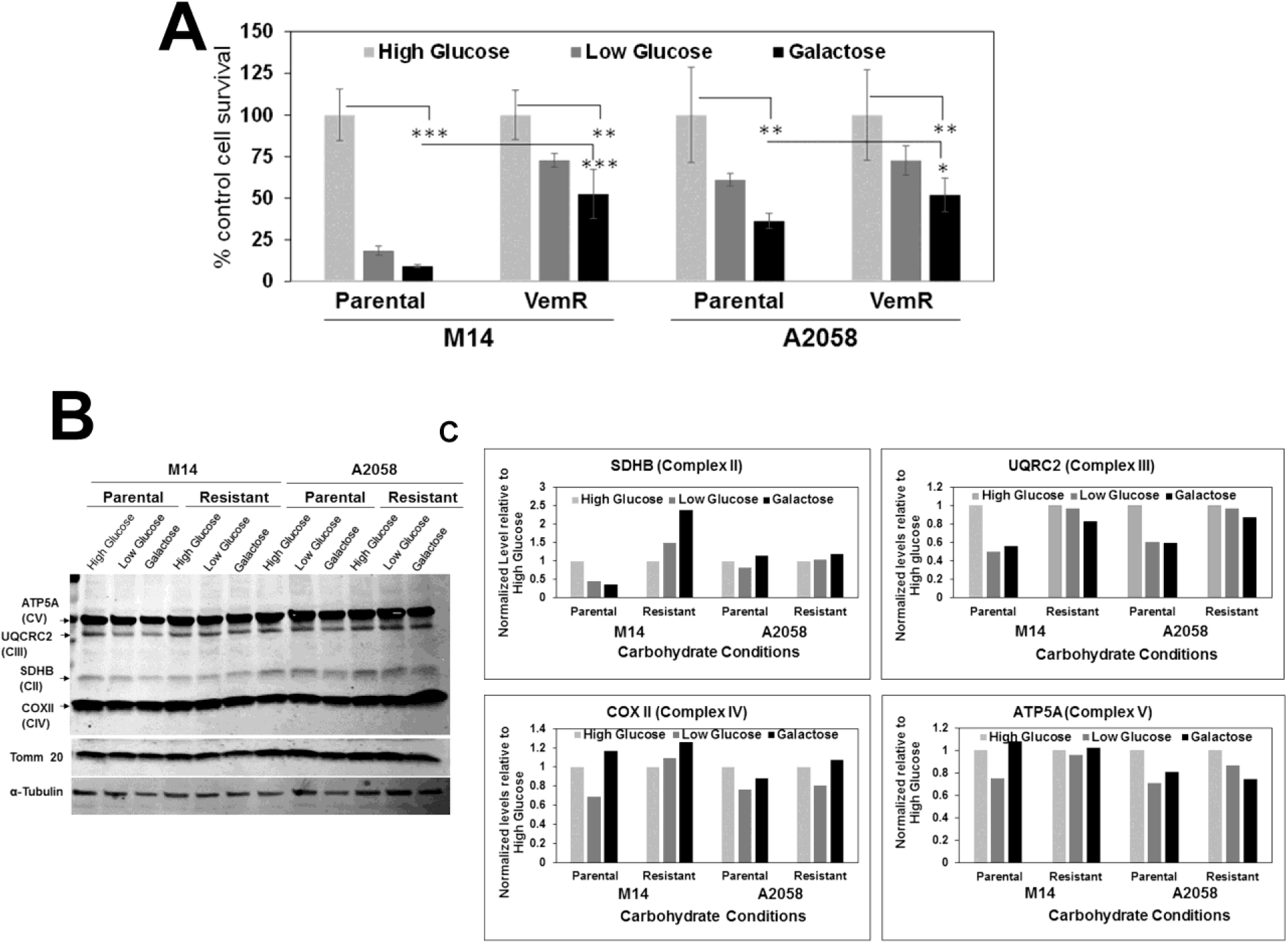
VemR melanoma cells display increased ability to utilize galactose as the energy source. (A) Survival analysis of parental and VemR M14 and A2058 cells under high glucose, low glucose or galactose conditions by MTT assays. Results are expressed as mean ± S.D. relative to cell survival in high glucose from three independent assays each performed in quadruplicates. *, P<0.05; **, P<0.01; ***, P<0. 001. (B) Western Blot analysis of OXPHOS complex proteins and (C) quantification of the levels of OXPHOS protein subunits normalized to Tomm20 and expressed relative to high glucose condition.

Next, we tested if the differences in glucose and galactose utilizations by parental vs. VemR isogenic melanoma models are associated with concomitant changes in OXPHOS complex II-V protein levels. Western blot analysis revealed that UQCRC2 decreased ∼50% in M14 and A2058 parental cells grown in galactose or low glucose media as compared to cells in high glucose medium whereas UQCRC2 levels remained elevated in VemR cells and were unaffected by galactose or low glucose (Fig. 6B, C). Levels of SDHB, a subunit of complex II, were also decreased in parental M14 cells under galactose or low glucose conditions and upregulated in their VemR counterpart (Figure 6B, C). No differences in SDHB levels were seen in A2058 derivatives, and the levels of COX II and ATP5A levels constituting complexes IV and V, respectively, were not impacted (Fig. 6B, C).

### β-catenin signaling promotes mitochondrial bioenergetics in VemR melanoma cells

Our data in Figures 1-3 showed that β-catenin signaling plays an important role in adaptive VemR development and that VemR development is associated with gain in mitochondrial bioenergetics. To determine whether β-catenin activity is involved in switch to mitochondrial metabolism, we analyzed the effects of ICG-001 on mitochondria activity. While vemurafenib treatment did not alter Mitotracker Red staining intensities or the percentage of cells with active mitochondria in M14 VemR or A2058 VemR cells, treatment with ICG-001 significantly decreased the percentage (P<0.01) and intensities of Mitotracker Red stained (P<0.001) cells (Fig. 7A-D). Immunofluorescence staining of β-catenin showed ICG-001 induced dramatic loss of intracellular β-catenin and relocation to the cell membrane in VemR cells (Fig. 7E, F). These data are consistent with the TOP/Flash reporter and β-catenin regulated gene expression data in Fig. 3E-G and confirm that ICG-001 induced sensitization of VemR melanoma cells results from ICG-001 induced inhibition of β-catenin activity.

**Figure 7.**
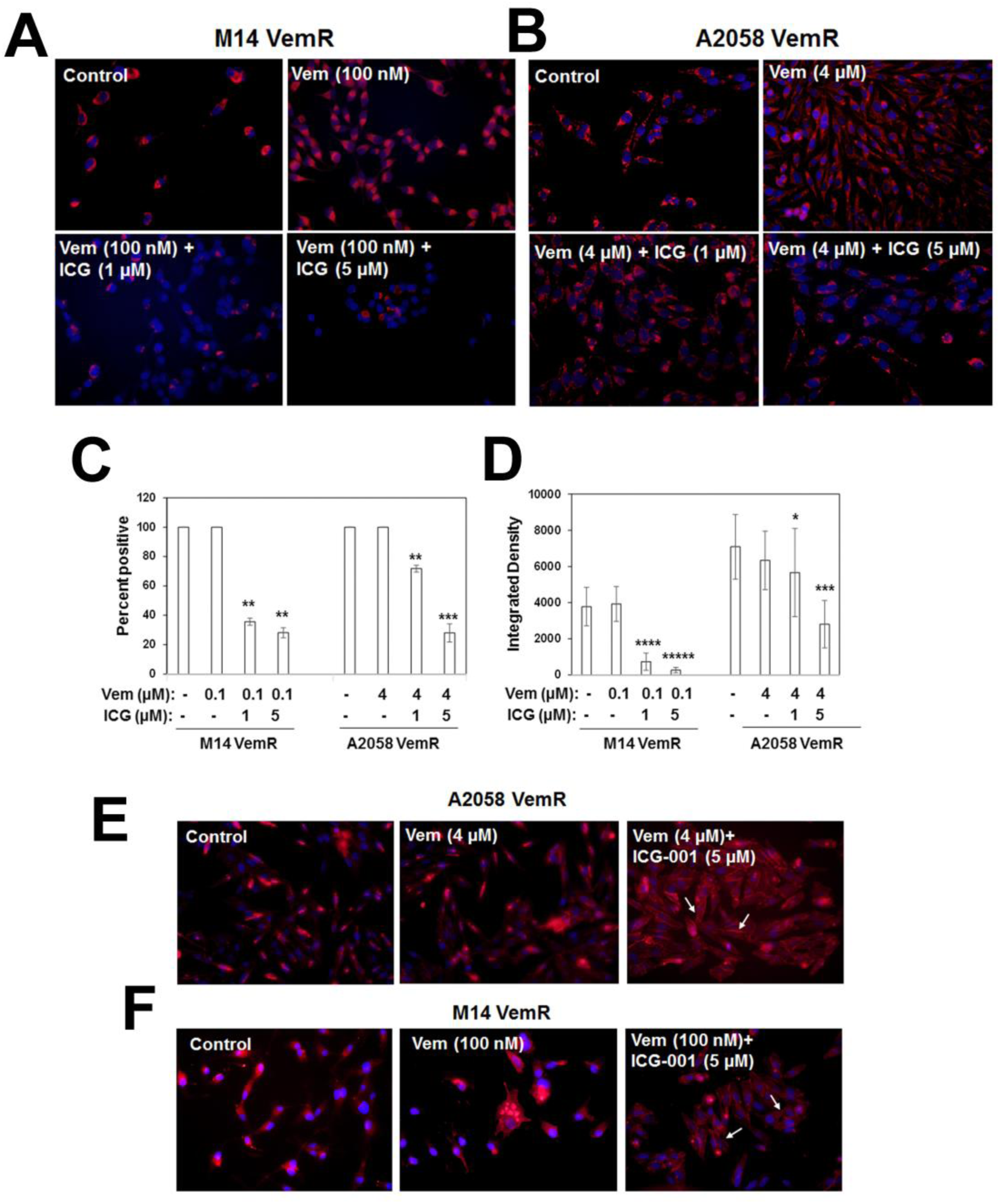
ICG-001 treatment decreases mitochondrial mass and activity in VemR melanoma cells. (A,B) MitoTracker DeepRed staining of VemR M14 and A2058 cells. Original magnification ×100. (C) Percent of MitoTracker Red positive cells and (D) active mitochondrial density determined using ImageJ. Approximately 30-50 cells from 3-5 fields were scored and results expressed mean ± S.D. *, P<0.05, **; P<0.01; ***, P<0. 001; ****, P<0.0001; *****, P<0.00001. (E,F) Immunofluorescence staining of β-catenin in VemR A2058 and M14 cells treated with ICG-001. Original magnification 40X. Arrows indicate ICG-001 induced decreases in intracellular β-catenin and its relocalization to the cell surface.

Cells cultured in galactose rely mostly on OXPHOS, which makes them more susceptible to mitochondrial toxicants.^37^ To determine if ICG-001 treatment will render different sensitivities to M14 or A2058 VemR cells grown under high glucose or galactose conditions, cells cultured in glucose or galactose media were treated with 0.5-10 µM ICG-001 and cell survival was analyzed by MTT assays. A dose-dependent reduction in cell survival was observed with increasing concentrations of ICG-001 under both high glucose and galactose conditions, suggesting involvement of β-catenin signaling in both glycolysis and OXPHOS pathways (Fig. 8A, B). To isolate the involvement of β-catenin signaling in OXPHOS activity, we measured OCR of M14 and A2058 VemR cells treated with ICG-001 under glucose and galactose conditions. ICG-001 treatment significantly decreased basal and maximal OCR in both A2058 VemR (P<0.001) and M14 VemR (basal, P<0.05; maximal, P<0.001) cells cultured in glucose media (Fig. 8C,E,G). ICG-001 treatment similarly reduced basal (P<0.001) and maximal (P<0.05) respirations of A2058 VemR cells cultured in galactose media (Fig. 8D, H), confirming involvement of β-catenin signaling in oxidative phosphorylation. M14 VemR cells grown in galactose or glucose media showed similar levels of basal OCR and were inhibited similarly by mitochondrial ATP synthase inhibitor oligomycin (Fig. 8E-H). These data suggest that M14 VemR cells have similar levels of basal OCR ascribed to ATP turnover under glucose and galactose conditions. However, in VemR cells cultured in galactose, injection of the uncoupler FCCP failed to stimulate the respiratory chain. It is interesting to note that while proton leak (or ion movements requiring proton motive force) and non-mitochondrial consumption of oxygen are decreased by ICG-001 under glucose in both M14 VemR and A2058 VemR cells (Fig. 8G), they were not affected by ICG-001 in M14 VemR and A2058 VemR cells cultured in galactose media (Fig. 8H). Since FCCP injection following oligomycin treatment fails to induce maximal respiration in cells grown under galactose, it suggests that respiration in these cells is controlled by oxidative phosphorylation. While ICG-001 treatment inhibited OCR in A2058 VemR cells (Fig. 8D, H), it had no effect on the bioenergetics of M14 VemR cells grown in galactose (Fig. 8F, H). These data suggest that β-catenin-dependent components for oxygen consumption are probably compromised under galactose. Analysis of ECAR presented as absolute rates from a stable baseline showed 5-10% increases in both VemR models cells as compared to their parental counterparts (Supplementary Fig. S4A, B). Treatment with ICG-001 induced 1.8- to 2-fold decrease in ECAR in M14 VemR and A2058 VemR cells, respectively, maintained in 25 mM glucose. The ECAR in 25 mM galactose medium was only ∼15-20% of what it was in 25 mM glucose medium, and while ICG-001 had no effect on ECAR in M14 VemR cells, it decreased ECAR in A2058 VemR cells cultured in galactose indicating continued maintenance of glycolytic metabolism and involvement of β-catenin pathway (Supplementary Fig. S4C, D).

**Figure 8.**
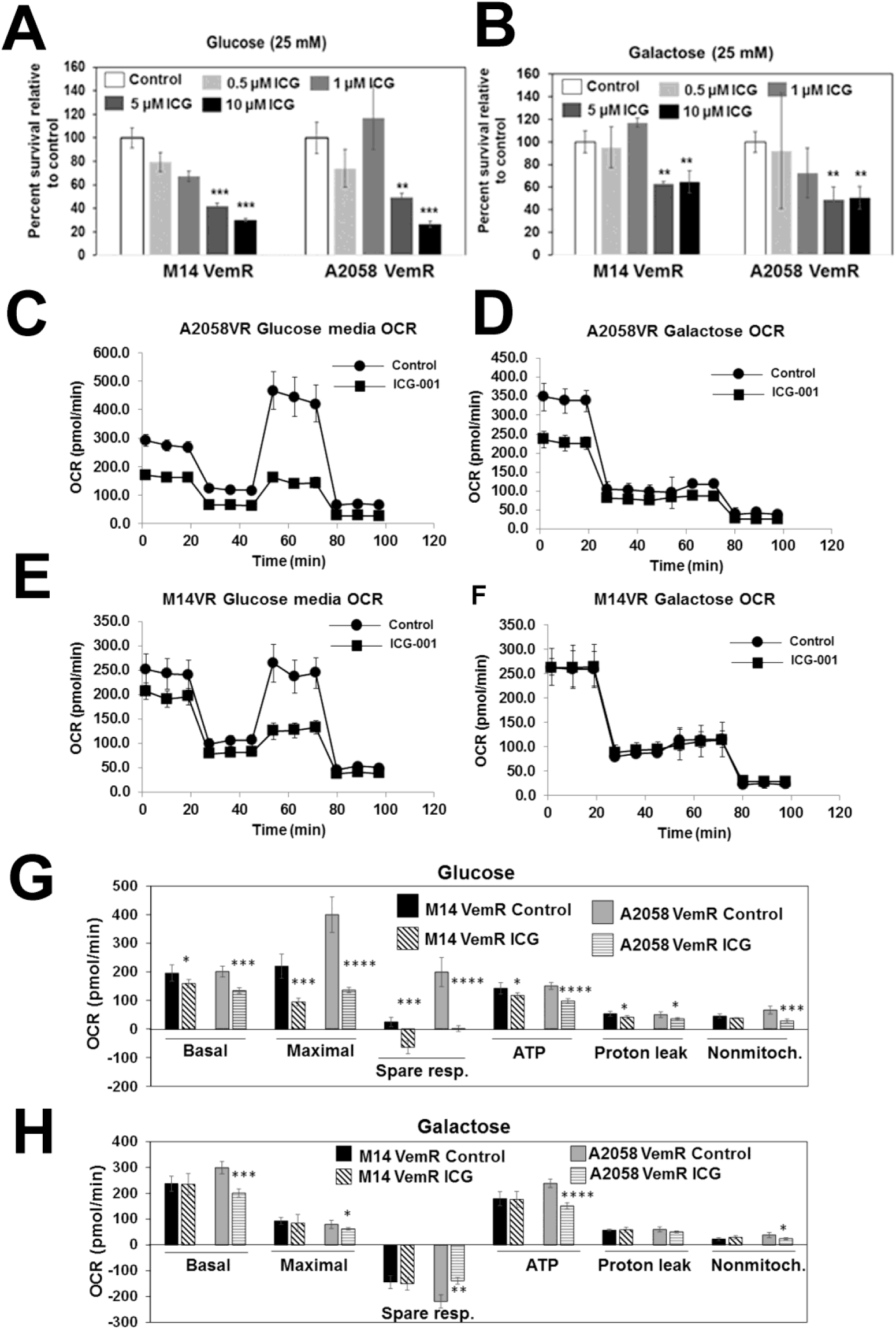
Vemurafenib resistant melanoma cells maintain sensitivity to ICG-001 under high glucose and galactose culture conditions. (A,B) Evaluation of ICG-001 sensitivities of VemR M14 and A2058 cells cultured in high glucose or galactose media using MTT assays. Results are expressed relative to controls mean ± S.D. from triplicates and two independent experiments. **, P<0.01, ***, P<0.001. (C-F) Bioenergetics analysis of VemR A2058 (C,D) and M14 (E,F) cells by Seahorse assay using mitochondrial stress kit under high glucose (C,E) and galactose (D,F). (G,H) Summary data calculated from the curves in C through F. *, P<0.05; **, P<0.01; ***, P<0. 001; ****, P<0.0001.

### Vemurafenib resistance development is associated with alterations in metabolic pathways involved with TCA cycle and pentose phosphate pathway

To gain an overview of the metabolic differences between parental and VemR melanoma cells, we performed a comparative analysis of metabolites using an LC-MS/MS-based targeted metabolomics platform that measures 254 metabolites involved in major human metabolic pathways. ^27^ The steady state levels of metabolites are shown in the heatmap (Fig. 9A). Pathway analysis of differential metabolites in parental vs. VemR M14 and A2058 cells revealed that metabolites involved in nucleotide (purine and pyrimidine biosynthesis), amino acid metabolism (alanine, aspartate and glutamate metabolism; arginine biosynthesis), and the citric acid (TCA) cycle were significantly impacted by VemR acquisition in both models (Fig. 9B, C). Metabolites involved in pentose phosphate pathway were strongly impacted (impact score = 0.5248, P = 0.0004) by adaptive VemR in the M14 model (Fig. 9B), whereas arginine/proline metabolism was significantly impacted (impact score = 0.51744, P = 0.003) in the A2058 model (Fig. 9C). Similar analysis of metabolites in control and vemurafenib treated melanoma PDXs Mel-14-108 (BRAF V600E) and Mel-14-089 (BRAF wild type) showed that vemurafenib treatment significantly impacted pathways related to glycine, serine and threonine metabolism, pentose phosphate pathway, NAD metabolism, pyruvate metabolism, arginine, glycolysis, and CoA biosynthesis only in BRAF mutant Mel-14-108 cells while vemurafenib had no effect on these processes in BRAF wild type Mel-14-089 cells (Supplementary Fig. S5 and Supplementary Tables S1A, B). These data confirm the selectivity of vemurafenib for mutant BRAF and mutant BRAF-driven metabolic pathways.

**Figure 9.**
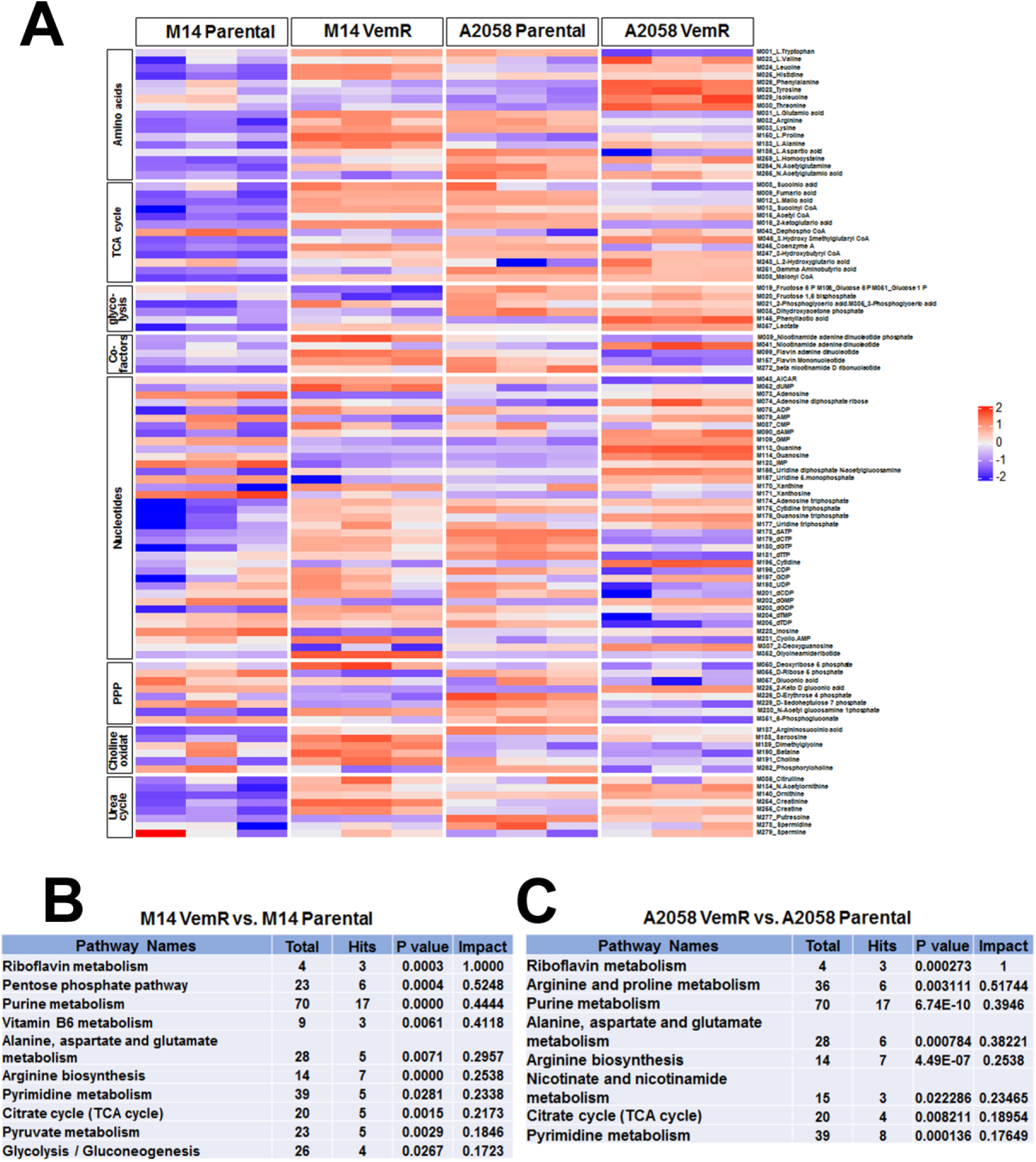
Acquisition of vemurafenib resistance alters the metabolome of melanoma cells. (A) Heatmap of targeted metabolomics data from isogenic parental and vemurafenib resistant melanoma cells. Each column represents a sample, and each row represents the expression levels of a single metabolite. The color scale of the heat map ranges from blue (low expression) to red (high expression). (B,C) Pathway analysis of top significantly altered metabolites and metabolic pathway impact between M14 VemR and parental (B), and A2058 VemR vs. parental cells were evaluated by Metabolite Set Enrichment Analysis (MSEA) using MetaboAnalyst. A cut-off value of 0.1 for pathway impact score was used consistently across multiple comparisons to filter less important pathways.

### VemR melanoma cells have increased ability to metabolize mitochondrial substrates to generate ATP

The metabolite analysis data in Figure 9 revealed an important role for mitochondria in VemR cells. To investigate the metabolic capacities and ATP generation, we assessed the mitochondrial functionality of parental and VemR counterparts using MitoPlate S-1 assay, which employs microplates pre-coated with various NADH and FADH2-producing metabolic substrates.^38,39^ We observed increases in metabolism of a variety of cytoplasmic (glycolytic), TCA cycle and other mitochondrial substrates by M14 VemR and A2058 VemR cells relative to their corresponding parental counterparts (Fig. 10A, B). However, A2058 VemR cells exhibited notably very high metabolic rates for TCA cycle substrates citric acid, isocitric acid, aconitinic acid, succinic acid, fumaric acid, malic acid, and α-keto-glutaric acid. The metabolic rate for these substrates was >2-3-fold higher in A2058 VemR cells compared to its parental control (Fig. 10B). The metabolic rates of other mitochondrial substrates such as β-hydroxy butyric acid, glutamine, and glutamic acid were also higher in A2058 VemR cells compared to their parental counterpart. Cytoplasmic substrates, such as glycogen, glucose 1-phosphate, glucose 6-phosphate, glycerol phosphate and lactic acid were also metabolized at higher rate in A2058 VemR cells compared to parental cells although the magnitude of metabolic rate was considerably less as compared to usage of TCA cycle substrates (Fig. 10B). Although the metabolic rates for TCA cycle and other mitochondrial substrates were ∼1.5-fold higher in M14 VemR cells compared to their parental cells, they were considerably less than in A2058 VemR cells (Fig. 10A). Most notably, the metabolic rates of key substrates of pentose phosphate pathway, gluconate-6-phosphate and glucose 6-phosphate were ∼3.6- and 15-fold, respectively, higher in M14 VemR cells compared to parental cells (Fig. 10A), and were more pronounced than in A2058 VemR cells (Fig. 10A). These data reveal enhanced metabolism via the mitochondrial and pentose phosphate pathways in adaptive VemR development. Since our data have shown an important role for β-catenin signaling in VemR development and metabolism, we determined the effect of ICG-001 on metabolism of cytoplasmic and mitochondrial substrates using MitoPlate S-1 assay. ICG-001 treatment selectively reduced the metabolic rates of D-gluconate-6-phosphate and D-glucose-6-phosphate by 2- and 9-fold, respectively, in M14 VemR cells as compared to parental counterparts while it had minimal effects on utilization of other cytoplasmic or mitochondrial substrates (Fig. 10C). Similar analysis of ICG-001 effects on A2058 VemR cells showed 1.5-2.8 decrease in metabolic rates of TCA cycle substrates citric acid, isocitric acid, aconitic acid, succinic acid, fumaric acid, malic acid and α-keto-glutaric acid but had little effect on D-gluconate-6-phosphate and D-glucose-6-phosphate (Fig. 10D). Taken together, the results from targeted metabolome and MitoPlate S-1 substrate utilization assays reveal a major role for β-catenin signaling in controlling TCA cycle (a central hub in mitochondrial OXPHOS) in A2058 VemR cells, and the pentose phosphate pathway in M14 VemR cells.

**Figure 10.**
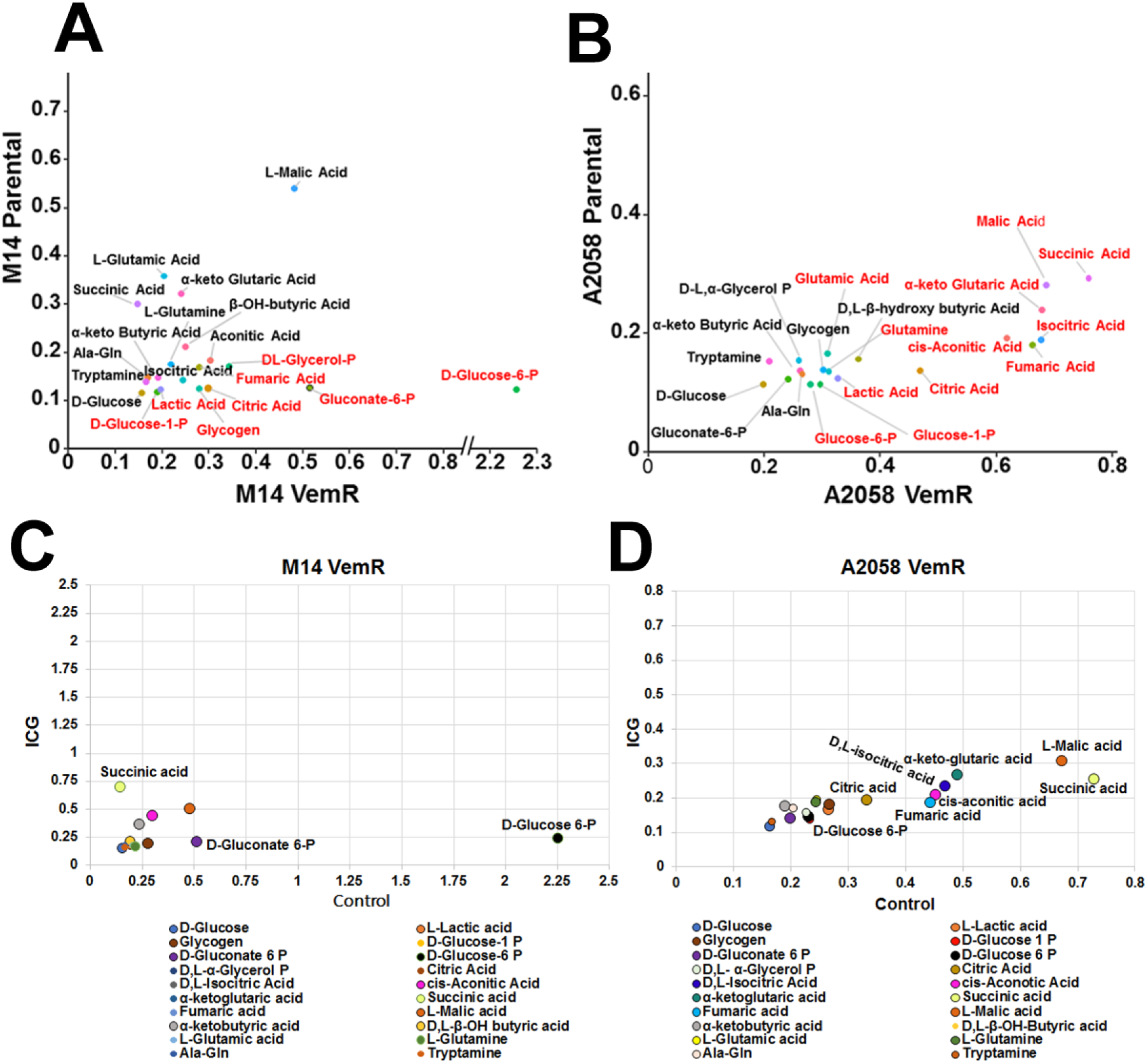
Mitochondrial metabolism and substrate preference analysis of vemurafenib resistant melanoma cells and ICG-001 inhibition. (A) Parental and VemR M14 cells or (B) parental and VemR A2058 cells were plated at a density of 30,000 cells/well in duplicates and the oxidation of a panel of cytoplasmic and mitochondrial substrates was assessed using Biolog MitoPlate-S1 assay. Maximal oxidation of each substrate was determined, and data are represented in scatter plots to show the maximal rates of various substrates between parental and resistant (A) M14 and (B) A2058 cell lines. Each dot represents a unique substrate. Substrates marked in red indicate those preferentially metabolized by the respective model. (C,D) Metabolism of cytoplasmic and mitochondrial substrates by M14 VemR (C) and A2058 VemR (D) cells treated with ICG-001. The data are represented in scatter plots to show the differences in maximal rates of substrate metabolism between control and ICG-001 treated VemR M14 and A2058 cells for the same substrate. Only substrates impacted by ICG-001 are indicated.

### Transcriptome analysis supports VemR associated alterations in β-catenin signaling and metabolic pathways

To analyze the molecular underpinnings of VemR, we characterized the global gene expression profiles of parental and VemR M14 and A2058 cells by whole transcriptome sequencing, and statistically significant differentially expressed genes (DEGs) were subjected to iPathway guide analysis to identify the affected processes. Volcano plot and meta-analysis using Venn diagram revealed 2158 and 2163 statistically significant DEGs (FDR ≤ 0.05) in M14 VemR vs. M14 and A2058 VemR vs A2058 cells, respectively, of which 867 genes were commonly expressed in M14 VemR and A2058 VemR groups (Supplementary Fig. S6A-E). Among the 867 genes, 282 and 200 genes, respectively, were upregulated and downregulated in both VemR models (Supplementary Fig. S6, D). Pathway analysis of the affected DEGs revealed enrichment of genes associated with 34 pathways in M14 VemR and A2058 VemR cells (Supplementary Fig. S6, E), and encompassed pathways associated with EGFR tyrosine kinase resistance, PPAR signaling, MAPK signaling, Ras signaling, cytokine receptor interaction, neuroactive ligand receptor interaction, and PI3K-Akt signaling pathways (Supplementary Fig. S6, F). Hierarchical cluster analysis showed clear separations of DEGs between parental and VemR M14 and A2058 groups (Fig. 11A, B), and identified upregulation of several genes associated with Wnt signaling in both VemR models (Fig. 11A, B). Among the Wnt/β-catenin regulated genes overexpressed in both VemR cells were TRPM1, ID3, ABCB5, TYRP1, PLA2G16 ^40^ and PHLDB2 ^41^ whereas SFRP1 and NDRG1,^42^ negative regulators of Wnt/β-catenin signaling were downregulated in both M14 VemR and A2058 VemR cells compared to their parental counterparts. ID3 (inhibitor of differentiation) and ABCB5 (subfamily B, member 5) transporter (a member of the ATP-binding cassette (ABC) superfamily) overexpression is reported in vemurafenib resistant melanomas.^43^ ABCB5 has been identified as a marker of melanoma-initiating cells^44^ and its expression is induced by β-catenin.^45^ Real-time RT-PCR analysis confirmed overexpression of ABCB5 in M14 VemR and A2058 VemR cells compared to their parental counterparts (Fig. 11C, D). Treatment with ICG-001 significantly decreased ABCB5 gene expression (P<0.01) and protein in both VemR models (Fig. 11E-G), confirming its regulation by the Wnt/β-catenin pathway. Expression of canonical Wnt signaling regulated genes MLANA, MITF, DCT, TYR and PMEL (premelanosome) were upregulated in A2058 VemR cells, however, these were downregulated in M14 VemR cells, suggesting potential reprogramming of pigmentation program in M14 cells upon acquisition of VemR (Fig. 11A, B). Pathway analysis of transcripts in BRAF mutant Mel-14-108 patient derived melanoma showed similar enrichment of pathways associated with cytokine-cytokine receptor interaction, ECM receptor interaction, and neuroactive ligand receptor interaction as in BRAF mutant M14 and A2058 models, whereas wild type BRAF Mel-14-089 patient derived melanoma cells showed enrichment of pathways associated with cell cycle, DNA replication, Fanconi anemia and DNA repair pathways, distinct from those regulated by mutant BRAF (Supplementary Table S2).

**Figure 11.**
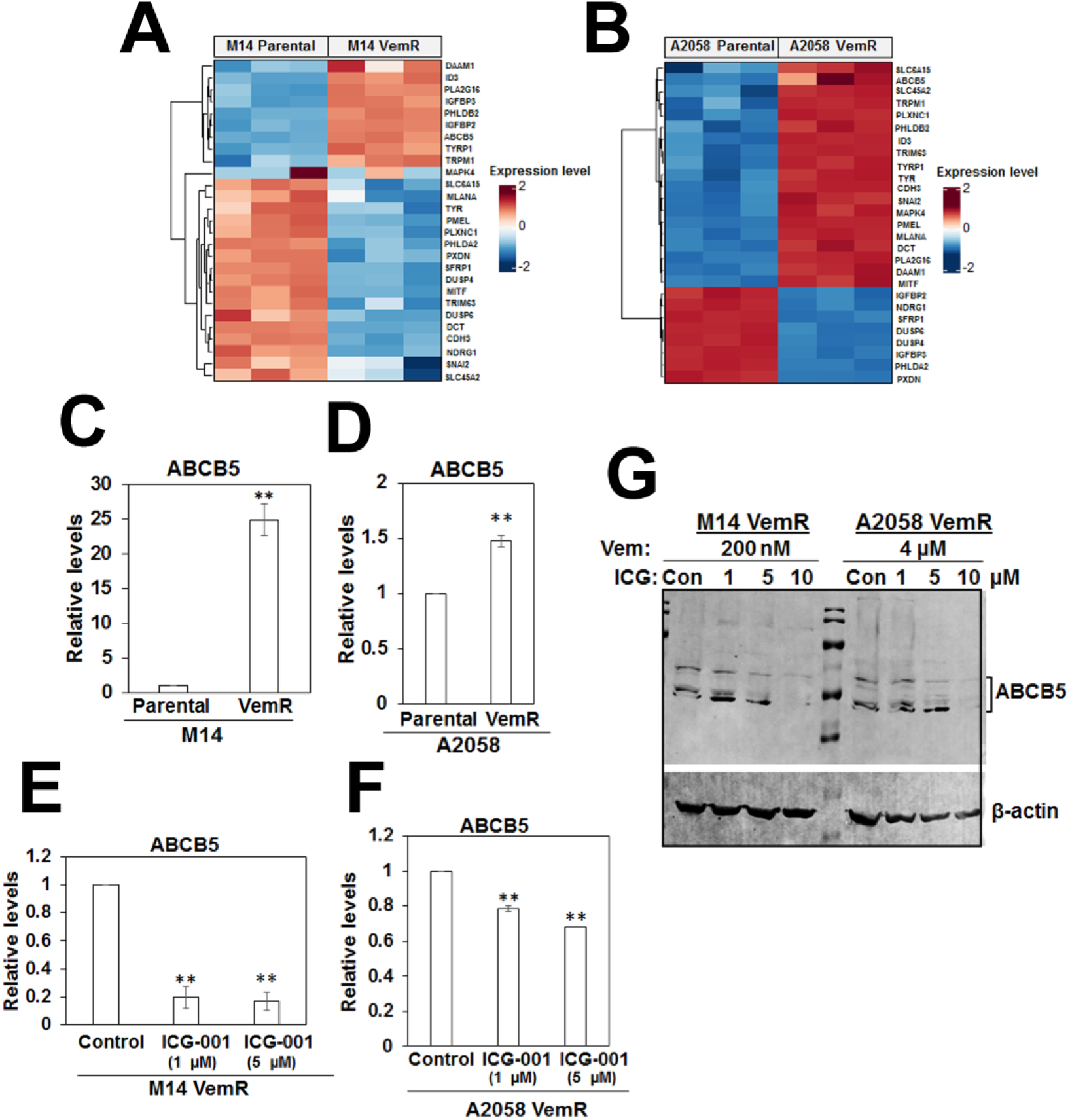
Canonical Wnt signaling regulated gene expressions are upregulated in vemurafenib resistant melanoma cells. (A,B) Heatmap of RNA-seq data comparing Wnt signaling regulated gene expressions in parental and VemR M14 (A) or parental and VemR A2058 cells. (C-F) Quantitative RT-PCR analysis of ABCB5 gene expression in parental and VemR M14 (C) and A2058 (D) cells, and regulation of ABCB5 expression by ICG-001 treatment in M14 VemR (E) and A2058 (F) cells. (G) Western blot analysis of ABCB5 and regulation by ICG-001 in VemR M14 and A2058 cells.

The relationship between vemurafenib resistance-impacted metabolic pathways and their regulatory mRNAs was analyzed in parental and VemR M14 and A2058 cells by querying the expressions of genes involved in the impacted metabolic pathways. Key pentose phosphate pathway (PPP) genes G6PD (a rate limiting enzyme in the PPP), RBKS (Ribokinase, a PfkB family member of carbohydrate kinases responsible for ribose phosphorylation) and PGM2 (involved in ribose and deoxyribose phosphate catabolism) were elevated in M14 VemR cells compared to their parental cells (Fig. 12A). In contrast, several genes associated with TCA cycle (CS, MDH1, OGDH, SUCLG1) were enriched in A2058 VemR cells compared to their parental counterparts (Fig. 12B). The higher expression of RBKS in M14 VemR cells is consistent with the elevated levels of 2-deoxy ribose 5-phosphate in M14 VemR cells (Fig. 9A). Overexpression of RBKS gene expression was validated by real-time RT-PCR analysis which showed 2.5-fold increase (P<0.001) in RBKS levels in M14 VemR cells compared to the parental cells (Fig. 12C). Treatment with ICG-001 significantly decreased RBKS gene expression in M14 VemR cells (P<0.0001), suggesting regulation by the Wnt/β-catenin pathway (Fig. 12E). RBKS gene expression was not significantly different between parental and VemR A2058 cells (Fig. 12D) and was unaffected by ICG-001 treatment (Fig. 12F). These data corroborate the metabolome (Fig. 10) and metabolite utilization data from MitoPlate S1 assay (Fig. 11) and indicate strong roles for TCA cycle and the pentose phosphate pathway in VemR development of A2058 and M14 cells, respectively. Taken together, our data suggest that acquisition of VemR and survival involve a key role for Wnt/β-catenin induced alterations in pathways associated with glucose catabolism.

**Figure 12.**
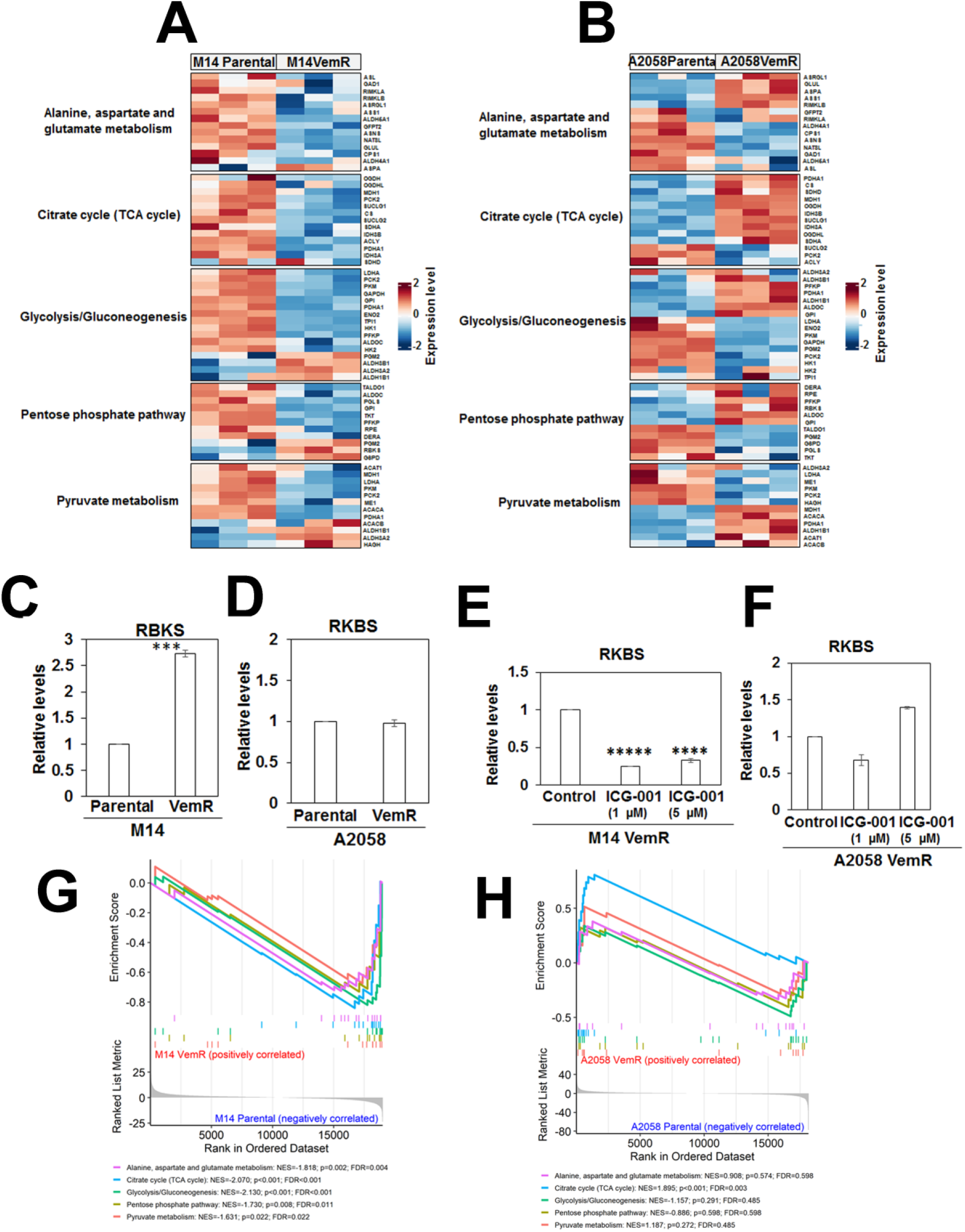
Transcript-metabolic pathway relationship. (A, B) Heatmap of genes regulating metabolic pathways impacted by VemR in M14 (A) and A2058 (B) isogenic pairs. (C-F) Quantitative RT-PCR analysis of RBKS gene expression in parental and VemR M14 (C) and A2058 (D) cells, and regulation of RBKS expression by ICG-001 treatment in M14 VemR (E) and A2058 (F) cells. ***, P<0. 001; ****, P<0.0001; *****, P<0.00001. (G,H) Gene set enrichment analysis (GSEA) plots of five representative pathway gene sets between M14 VemR and M14 parental groups (G), and (H) between A2058 VemR and A2058 parental cells.

Gene set enrichment analysis (GSEA) was conducted on genes corresponding to five metabolic pathways (Fig. 12G, H). All five pathways were significantly enriched in the M14 parental group, with negative normalized enrichment scores (NES) indicating downregulation in M14 VemR relative to the parental counterpart. Glycolysis/gluconeogenesis (NES = -2.130, p < 0.001, FDR < 0.001) and citrate cycle (TCA cycle) (NES = -2.070, p < 0.001, FDR < 0.001) pathways showed the highest negative NES values, followed by alanine, aspartate, and glutamate metabolism (NES = -1.818, p = 0.002, FDR = 0.004), pentose phosphate pathway (NES = -1.730, p = 0.008, FDR = 0.011), and pyruvate metabolism (NES = -1.631, p = 0.022, FDR = 0.022) (Fig. 12G). The citrate cycle pathway was most enriched in the A2058 VemR cells compared to its parental counterpart, with a positive NES (NES = 1.895, p < 0.001, FDR = 0.003) (Fig. 12H).

## Discussion

The aim of this work was to get a detailed understanding of the mechanisms and metabolic choices involved in development of adaptive VemR by integrating metabolome and transcriptome data from isogenic models of VemR metastatic melanoma and patient derived brain metastatic melanomas. For this study, we developed two isogenic human metastatic melanoma models of VemR using M14 and A2058 cells that carry *BRAFV600E* mutation, and included two patient derived brain metastatic melanomas with wild type (Mel-14-089) or V600E mutant (Mel-14-108) BRAF. Our results show that canonical Wnt/β-catenin signaling plays a major role in metabolic switch underlying adaptive VemR and contributes to increased mitochondrial mass and bioenergetics. Treatment with MEK (U0126 or PD098059) or PI3K (LY294002) inhibitors only partially suppressed proliferation of VemR cells, indicating that our VemR cells were also cross-resistant to MEKi and PI3Ki. Since inhibition of β-catenin transcriptional activity with ICG-001 decreased β-catenin regulated gene expression, inhibited ERK1/2 phosphorylation, and decreased mitochondrial activity and respiration with restoration of vemurafenib sensitivity, our data reveal an important role for β-catenin crosstalk with BRAF and ERK pathways in VemR development. These findings are consistent with previous reports linking activation of β-catenin signaling with VemR. ^33,34^

The hallmark of metabolic reprogramming by cancer cells is a shift towards glycolysis or Warburg effect as the key energy production mechanism,^46^ and a trend towards elevated mitochondrial biogenesis and oxidative phosphorylation has been observed in BRAF^V600E^ melanomas that develop drug resistance.^12,47^ Metabolic reprogramming is regulated by both canonical and noncanonical Wnt signaling ^48^ in many tumor types including melanoma, ^49^ and Wnt signaling regulates Warburg effect by inducing expression of key regulators of glycolysis.^50^ Studies suggest an important role for PTEN function in mediating Wnt/β-catenin induced upregulation of mitochondrial dynamics.^51,52^ In this context, it is interesting to note that despite differences in PTEN status, both PTEN wild type M14 and PTEN null A2058 VemR cells exhibit increased mitochondrial activity, bioenergetics and Wnt/β-catenin activity, and treatment with the Wnt/β-catenin antagonist ICG-001 similarly reduces mitochondrial activity and OCR in both VemR models, suggesting a PTEN-independent role of Wnt/β-catenin signaling in mitochondrial regulation of VemR.

Mitochondrial stress assays using Seahorse showed that although M14 VemR cells have significantly lower basal and maximal OCR compared to their parental counterparts, their spare or reserve respiratory capacity is slightly higher than their sensitive counterparts. The spare respiratory capacities of A2058 VemR cells are also higher than their corresponding parental counterparts, and analysis of OXPHOS levels showed upregulation of UQCRC2, a component of complex III, in both resistant models. These data provide support for a common role for mitochondria in coping with drug induced metabolic stress. To verify switch to mitochondrial metabolism, the OCRs of VemR cells were compared under glucose and galactose conditions since galactose increases a cell’s dependence on mitochondrial ATP production. ^53,54^ Similar levels of basal OCR were seen under both conditions suggesting efficient usage of the mitochondria by both A2058 and M14 VemR cells for ATP production. Treatment with the β-catenin inhibitor ICG-001 dramatically reduced basal and spare respiratory capacities, and ATP levels under both glucose and galactose conditions in A2058 VemR cells. However, this inhibition occurred only under glucose culture conditions in M14 VemR cells as ICG-001 treatment had no effect on oxygen consumption when M14 VemR cells were cultured in galactose or forced to use mitochondria. These data reveal differences in Wnt/β-catenin pathway dependence with A2058 VemR cells exhibiting greater reliance on the Wnt/β-catenin pathway for OXPHOS compared to M14 VemR cells. Both isogenic models of VemR have slightly higher or maintain similar levels of ECAR compared to their sensitive counterparts indicating that the glycolytic flux of VemR cells remains stable despite switch to mitochondrial metabolism. Although galactose itself should not inhibit glycolysis flux, culturing in galactose media resulted in ∼5-fold decrease in ECAR compared to cells in glucose. The reason for this is unclear, however, it is possible that the forced reliance on mitochondrial ATP production could redirect pyruvate generated by the glycolysis flux away from lactic acid production and into the mitochondria for OXPHOS. In such a case, this should result in reduction in lactate production. On the contrary, our data from targeted metabolite analysis and MitoPlate S-1 assays show higher levels of lactate, suggesting either diversion of galactose-derived carbon away from glycolysis and into the pentose phosphate pathway (PPP) ^55^ or a more complex OCR-ECAR relationship under galactose treatment. Treatment with β-catenin inhibitor reduced ECAR under both sugar conditions in A2058 VemR cells and only under glucose in M14 VemR cells, indicating dual roles for Wnt/β-catenin pathway in modulating glycolytic and OXPHOS metabolism.

RNA-seq analysis revealed numerous genes regulated by Wnt/β-catenin pathway (ABCB5, TYRP1, ID3, TRPM1) were upregulated while the negative Wnt/β-catenin regulator SFRP1 was downregulated in both M14 and A2058 VemR cells as compared to their parental counterparts. Studies have reported upregulation of ID3 (inhibitor of differentiation 3, a helix-loop-helix TF superfamily member) in melanomas of patients treated with a BRAFi and silencing ID3 resulted in enhanced sensitization of melanoma to vemurafenib.^56^ RNA-seq analysis showed MITF (a major regulator of mitochondrial biogenesis) gene overexpression in A2058 VemR cells but not in M14 VemR cells although MITF protein expression was induced, suggesting either low MITF transcription rates or transcript stability. In this regard, it is interesting to note that MITF inhibitor ML329 similarly inhibited proliferation of both VemR models supporting its role in mitochondrial activity (Supplementary Fig. S2). RNA-seq analysis (and further validated by qRT-PCR and protein analysis) also identified overexpression of ABCB5, a member of the ATP-binding cassette (ABC) transporter superfamily, in both VemR models. ABC transporters facilitate the ATP-driven efflux of drugs across cellular membranes,^57,58^ and is associated with multi-drug resistance in several cancers including melanoma.^59^ shRNA-mediated silencing of ABCB5 increased sensitivities of human colorectal cancer cells to 5-FU.^60^ ABCB5 is a marker of melanoma aggressiveness, multidrug resistance and stemness,^40,61,62^ and has been reported to induce unique modifications of metabolism in melanoma-initiating cell lines.^63^ The ABCB5 gene encodes several isoforms including ABCG5FL, a full transporter, ABCB5β, a half transporter, and heterodimers of ABCB5β with ATPase activity.^64^ In melanoma, its expression is activated by Wnt/β-catenin and MITF, ^46^ and as expected ICG-001 treatment suppressed ABCB5 mRNA as well as expressions of ABCB5 immunoreactive isoforms in both our VemR models.

Metabolomics and mitochondria-function based MitoPlate S-1 assays identified upregulation of TCA cycle metabolism in both M14 and A2058 VemR cells. However, the metabolic rates of TCA cycle substrate utilization as measured by MitoPlate S-1 assays were considerably higher in A2058 VemR cells as compared to M14 VemR cells, and suggest that the net effects of these genes could lead to hyperactivation of the TCA cycle. Although both M14 VemR and A2058 VemR cells undergo mitochondrial metabolic reprogramming, the magnitude of metabolic shifts differs greatly and appears to correlate with the vemurafenib tolerance thresholds of individual melanomas as melanomas with greater levels of de novo or acquired drug tolerance (such as A2058) are better at exploiting mitochondrial pathways for their bioenergetic needs. Our data from metabolomic analysis of patient derived metastatic melanoma Mel-14-108 further confirmed the selectivity of vemurafenib for BRAF mutation as none of the pathways (glycine/serine/threonine metabolism, PPP, nicotinamide metabolism, pyruvate metabolism, glycolysis) targeted by vemurafenib in vemurafenib sensitive Mel-14-108 cells were affected by vemurafenib in wild type BRAF Mel-14-089 cells (Supplementary Fig. 5 and Supplementary Table 1A,B). Consistent with the increased reliance of A2058 VemR cells on the mitochondria for energy production, integration of metabolomics pathway analysis data with RNA-seq analysis confirmed overexpression of several genes associated with the TCA cycle (CS, PDHA1, MDH1, OGDH, IDH3B, IDH3A, SUCLG1, OGDHL) in A2058 VemR cells. The two most important products produced from the pentose phosphate pathway (PPP) branch are ribose 5-phosphate used to make DNA and RNA, and the antioxidant nicotinamide adenine dinucleotide phosphate (NADPH) for nucleotide synthesis and maintenance of redox homeostasis^65^ for enabling cancer cells survive drug induced metabolic stress.^66^ Our results show that among the genes associated with PPP (G6PD, a gatekeeper and rate-limiting enzyme in the PPP; RBKS, critical for synthesis of ribose 5-phosphate; and PGM2, a regulator of ribose and deoxyribose phosphate catabolism) ^67,68^ are increased in M14 VemR cells, and are consistent with the robust utilization of PPP substrates glucose 6-phosphate and gluconate 6-phosphate, and increases in levels of deoxy ribose 5-phosphate. MitoPlate S-1 assays conducted with A2058 and M14 VemR cells treated with β-catenin antagonist showed suppression of TCA cycle and PPP substrate utilizations in A2058 VemR and M14 VemR cells, respectively. An important role for Wnt/β-catenin in inducing G6PD expression and activation of PPP and conferring chemoresistance is reported.^69^ Real-time RT-PCR analysis of RBKS confirmed its overexpression in M14 VemR cells and consistent with MitoPlate S-1 data, RBKS gene expression is inhibited by ICG-001. These results implicate an important role for Wnt/β-catenin pathway in modulating mitochondrial and PPP metabolism in VemR cells, and indicate that the magnitude of shift towards mitochondrial metabolism is higher in melanoma cells with greater vemurafenib tolerance thresholds while melanoma cells with lower tolerance limits divert metabolism to the PPP for nucleotide synthesis and redox regulation.

In summary, our results show that Wnt/β-catenin signaling plays a major role in adaptive VemR by coordinating the expression of critical genes involved in controlling melanoma metabolism and drug resistance, and that targeting this signaling could produce strong interference of these activities in BRAFi, MEKi and PI3Ki resistant melanomas and enhance vemurafenib sensitivity. Our data also show for the first time that the extent and duration of metabolic reprogramming is defined by vemurafenib tolerance limits of individual melanomas as melanomas with elevated thresholds of either intrinsic or acquired drug tolerance adeptly exploit mitochondrial pathways for their bioenergetic needs whereas melanomas with low drug tolerance thresholds switch to PPP, an alternative pathway of glycolysis, for survival.

## Data availability statement

The datasets generated and analyzed for this study can be found in the gene expression omnibus (GEO) (http://ncbi.nlm.nih.gov/geo/) database through accession # GSE284250.

## Author contributions

MPS conceived, designed, analyzed the results and directed the project. PNM and MA conducted the experiments and analyzed data, SA and NP conducted and analyzed the Seahorse flux assays, JL and SK assisted with metabolomic analysis, and SK, KG and HJ assisted with RNA-seq data analysis. MPS and PNM wrote the original draft, and all authors reviewed and edited.

## Funding

This work was supported by 3 Balls Racing (to MPS), R21CA288314-01 (to MPS), GrantBoost from the OVPR of Wayne State University (to MPS) and Molecular Therapeutics Program of Karmanos Cancer Institute (to MPS). The Genome Sciences Core, Biostatistics and Bioinformatics Core, and Pharmacology Core facilities are supported by NIH Center grant P30 CA022453 to the Karmanos Cancer Institute at Wayne State University.

## Conflict of interest

The authors declare that the research was conducted in the absence of any commercial or financial relationships that could be construed as a potential conflict of interest.

## Notes

### Competing Interest Statement

The authors have declared no competing interest.

### Summary of Updates

No changes were made to the original version.

