## Supplementary Data for "Relationship between melanoma vemurafenib tolerance thresholds and metabolic pathway choice and Wnt signaling involvement"

**Figure S1. Sensitivity analysis of BRAF mutant human metastatic melanoma cells to vemurafenib.** (A,B) MTT assays of M14 and A2058 cells. Results are expressed as mean ± S.D. (percent of control cell survival) from three independent experiments. (C,D) Clonogenic analysis of M14 and A2058 cells. (E,F) Visual depiction of representative wells captured using GelCount Oxford Optronix. Results are expressed as mean ± S.D. (percent of control colony formation efficiency) from three independent experiments. *P<0.05; **P<0.01; ***P<0.001; ****P<0.0001.

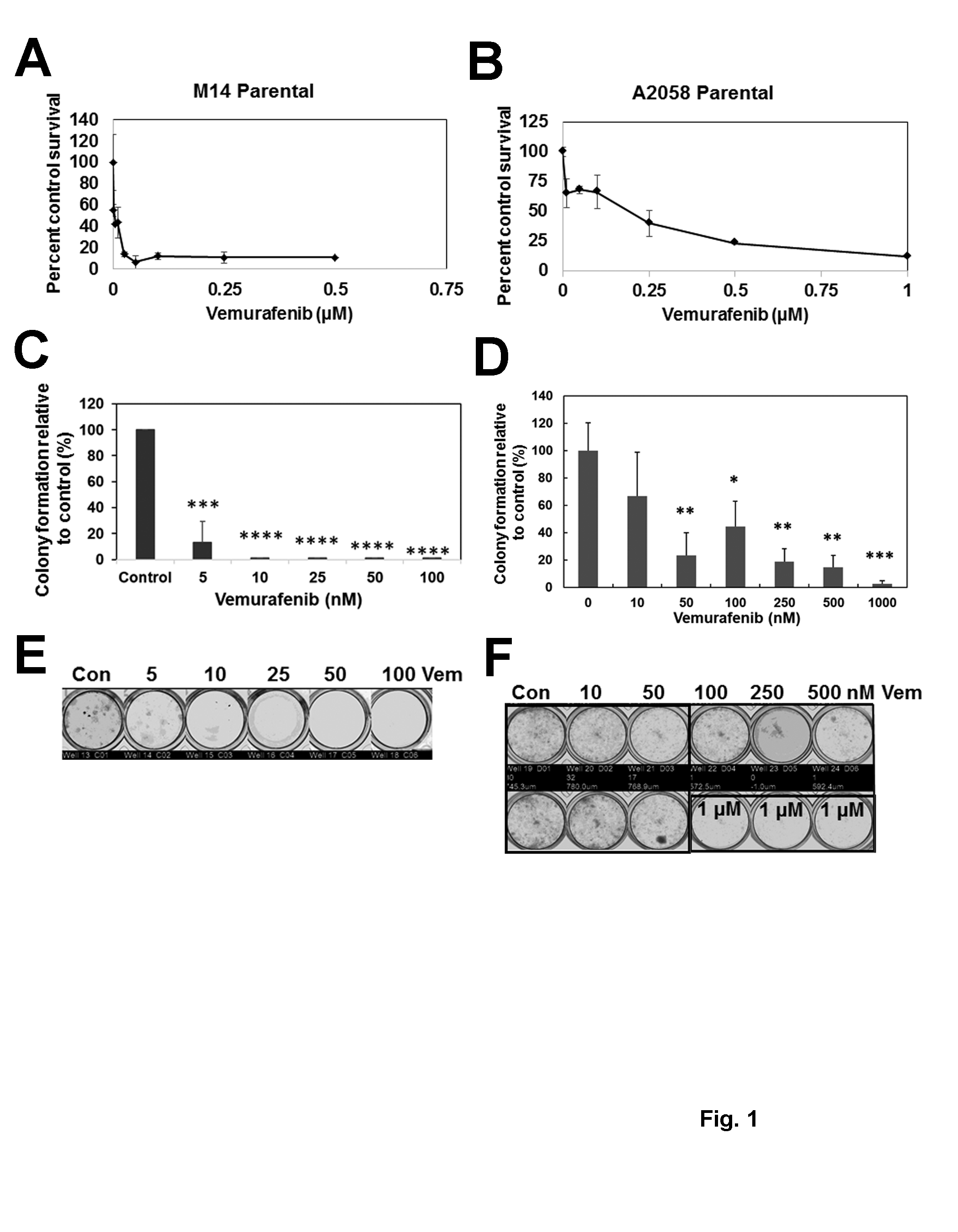

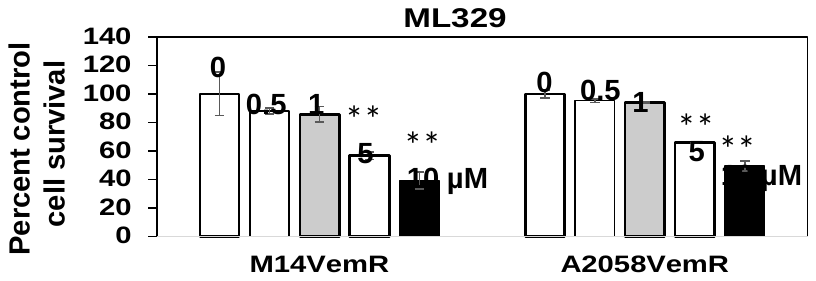
**Figure S2. Sensitivity analysis of VemR melanoma cells to MITF inhibitor ML329.** Cells were treated with the indicated concentrations of ML329.

**Figure S3. OXPHOS protein analysis in parental and VemR melanoma cells.** (A, B) Western blot analysis of subunits of OXPHOS complexes in M14 and A2058 isogenic pairs treated with the indicated concentrations of vemurafenib. Quantification of the levels of OXPHOS protein subunits normalized to Tomm20 and expressed relative to control in M14 (C-I) and A2058 (D-J) isogenic cells.

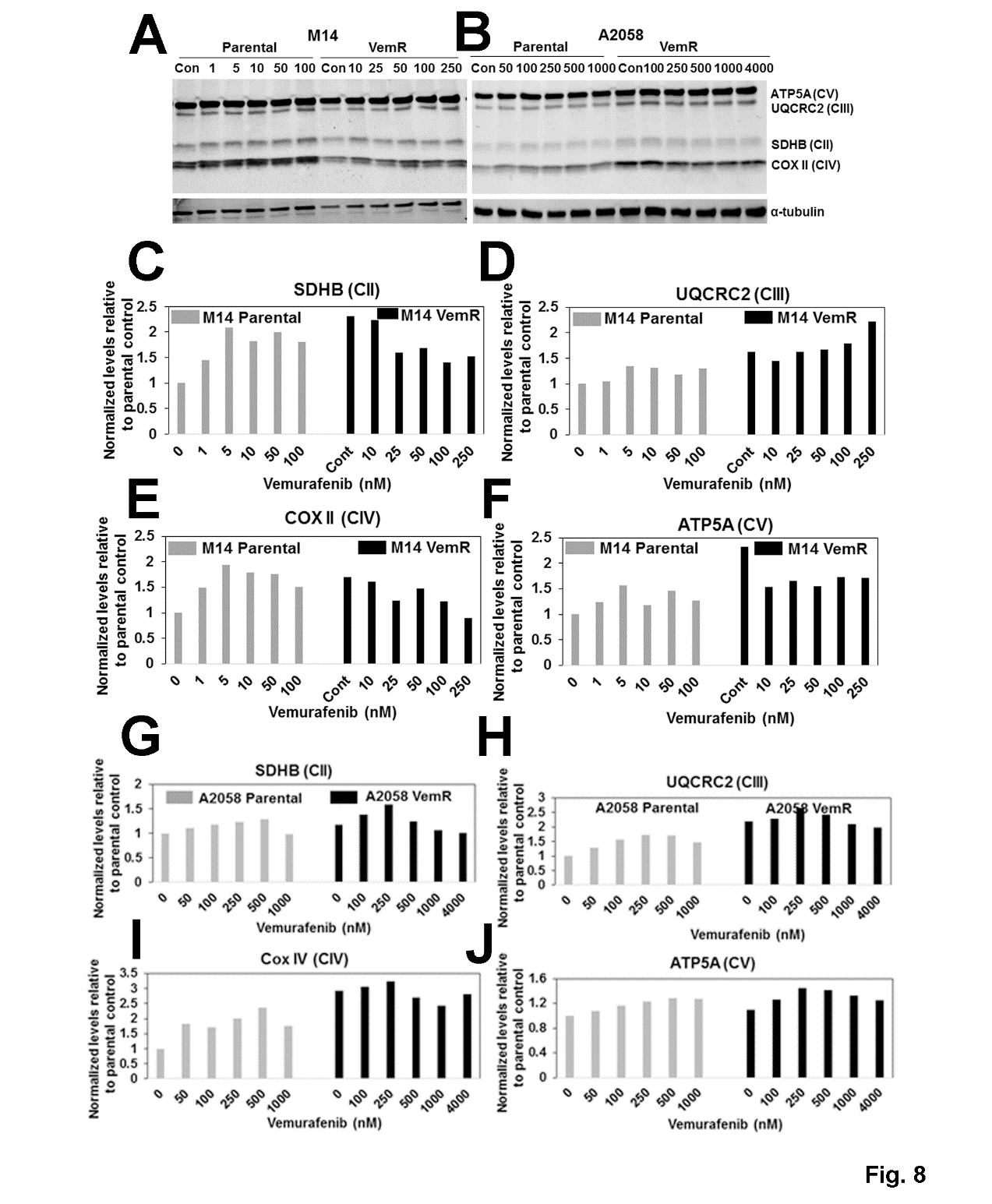

**Figure S4. ECAR rates of parental and VemR M14 and A2058 cells and effect of ICG-001 under glucose and galactose conditions. (A) Comparisons of ECAR rates of isogenic M14 and A2058 parental and VemR cells. (B) ICG-001 effects on ECAR under glucose and galactose conditions.**

**A**

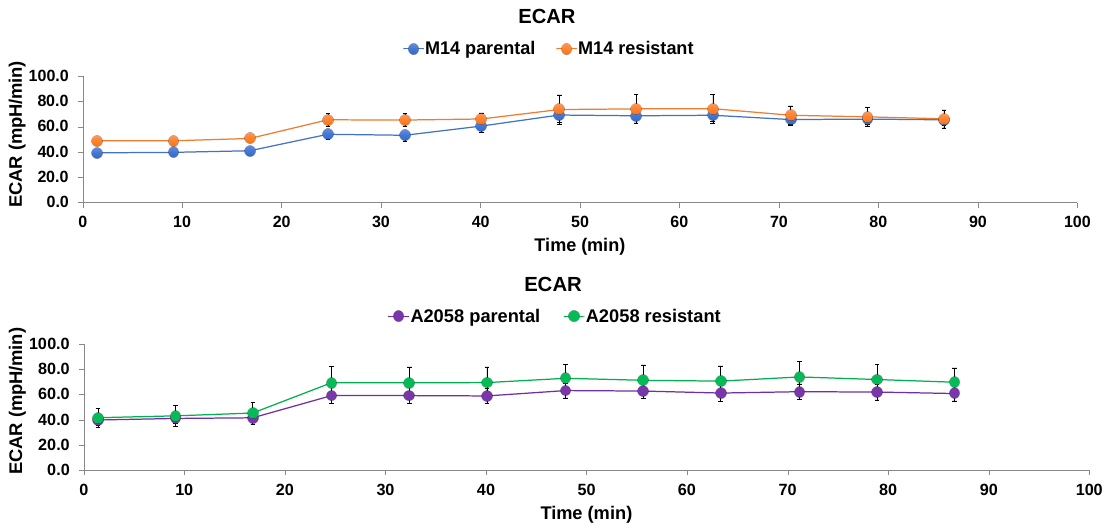

**B**

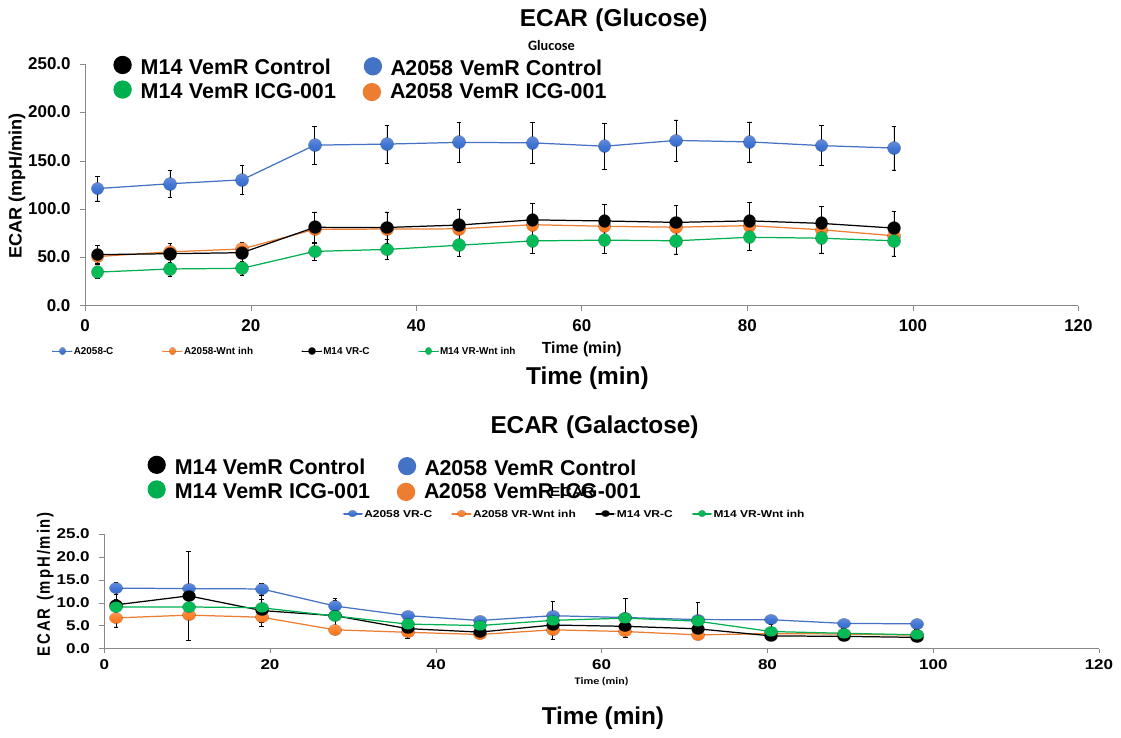

**Figure S5. Impact of vemurafenib on metabolome of melanoma patient derived cells Mel-14-108 (BRAF mutant) and Mel-14-089 (wild type BRAF).** .

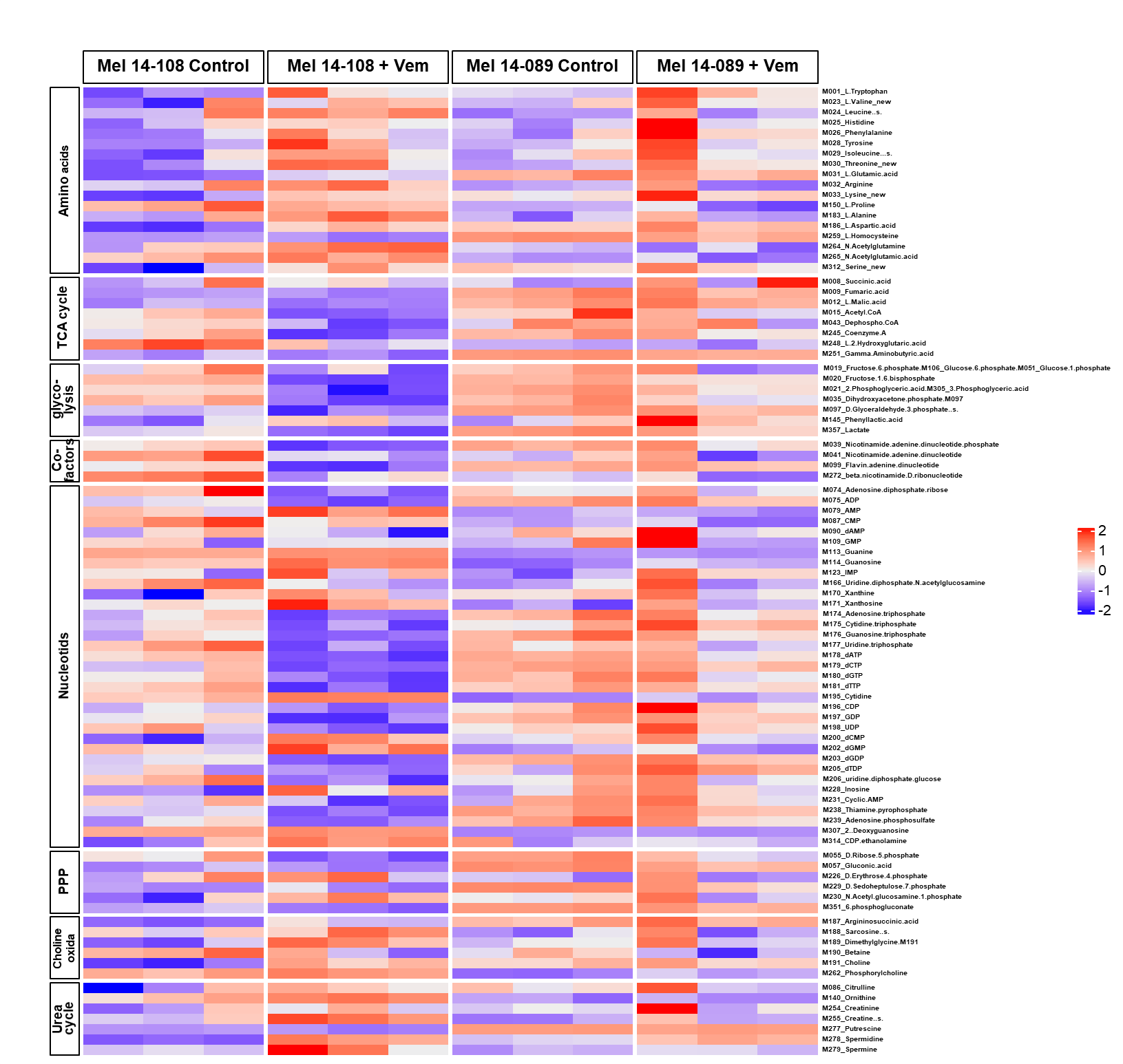

**Figure S6. Vemurafenib resistance acquisition alters the transcriptome of melanoma cells.** (A,B) Volcano plots of RNA-seq data comparing VemR vs. parental M14 (A) or VemR vs. parental A2058 (B) cells. Total number and differentially expressed genes (DEGs) genes 14756 and 3025, respectively (M14 VemR vs. M14 Parental), and 14079 and 3030, respectively (A2058 VemR vs. A2058 Parental) measured at 5% FDR and fold-change (FC) of ≥2. (C) Venn diagram of DEGs and (D) directional distribution of DEGs in M14 VemR vs. parental, and A2058 VemR vs. parental cells. (E) Meta analysis of pathways enriched in VemR M14 and A2058 cells, and (F) VemR impacted canonical pathways identified by iPathway Guide.

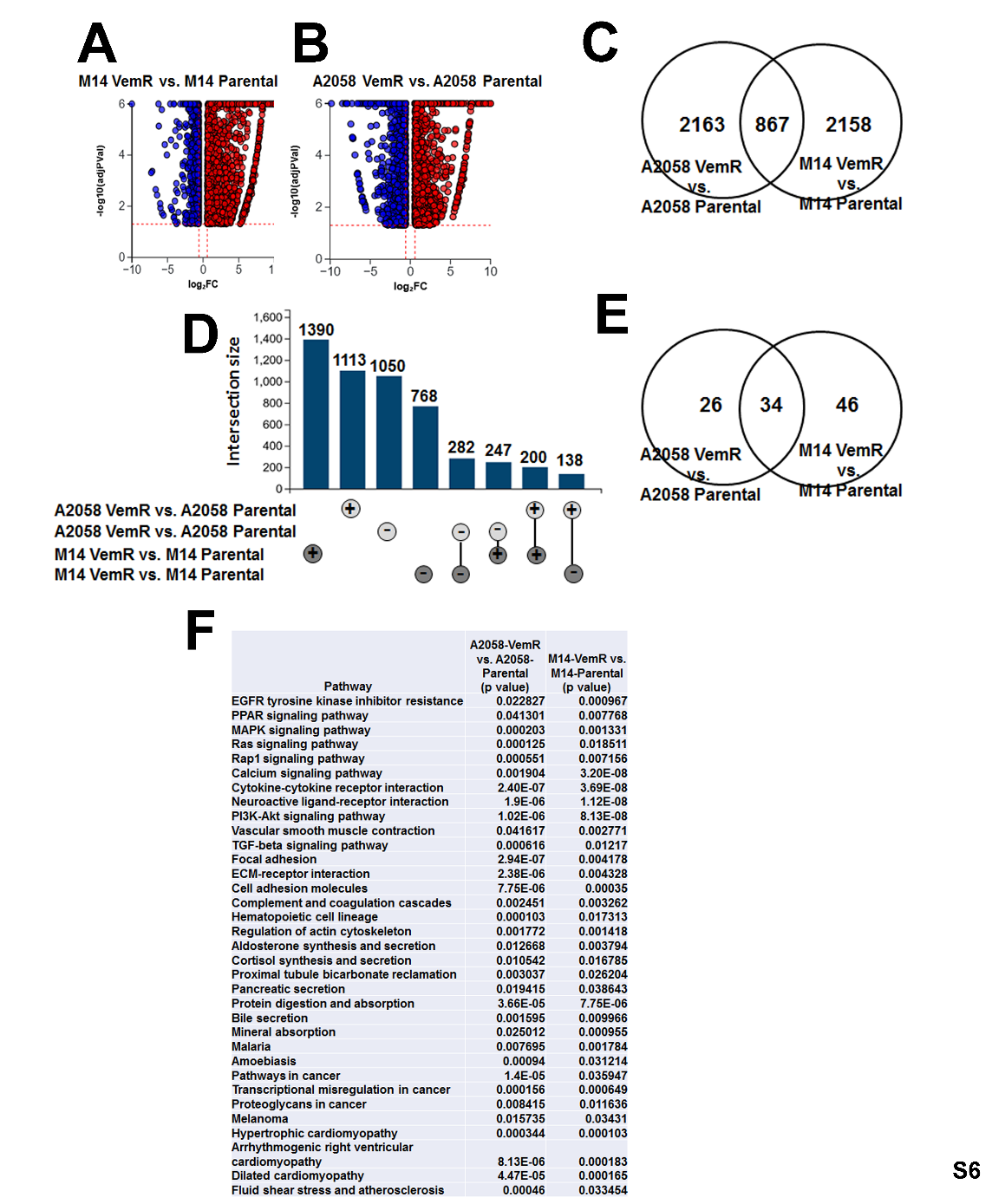

**Table S1. Metabolic pathways impacted by vemurafenib in melanoma patient derived cells expressing mutant BRFAF (Mel-14-108) or wild type BRAF (Mel-14-089).**

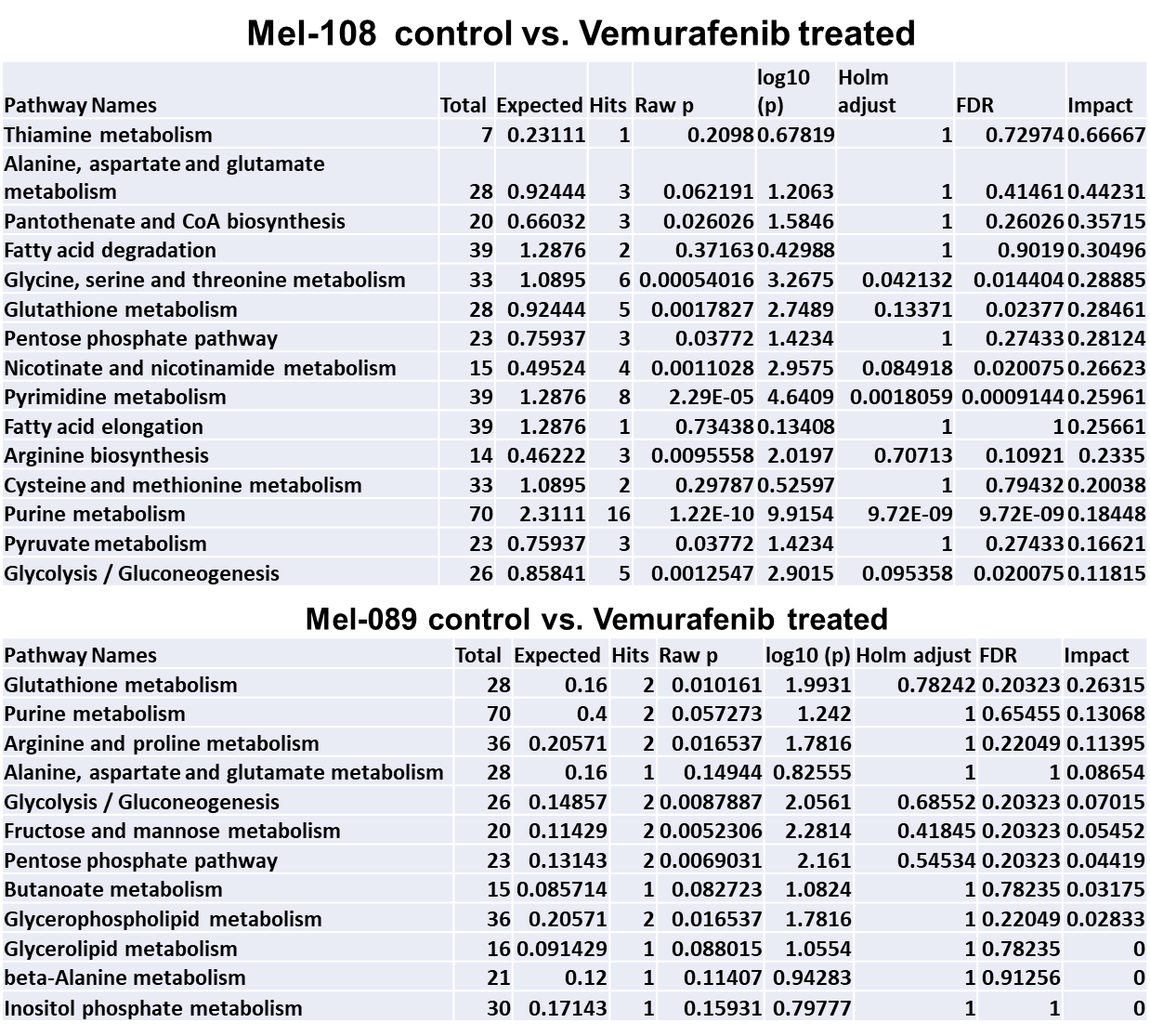

**Table S2. Pathway analysis of transcripts show distinct enrichment of pathways in patient derived melanoma cells with mutant BRAF (Mel-14-108) vs. wild type BRAF (Mel-14-089) cells.**

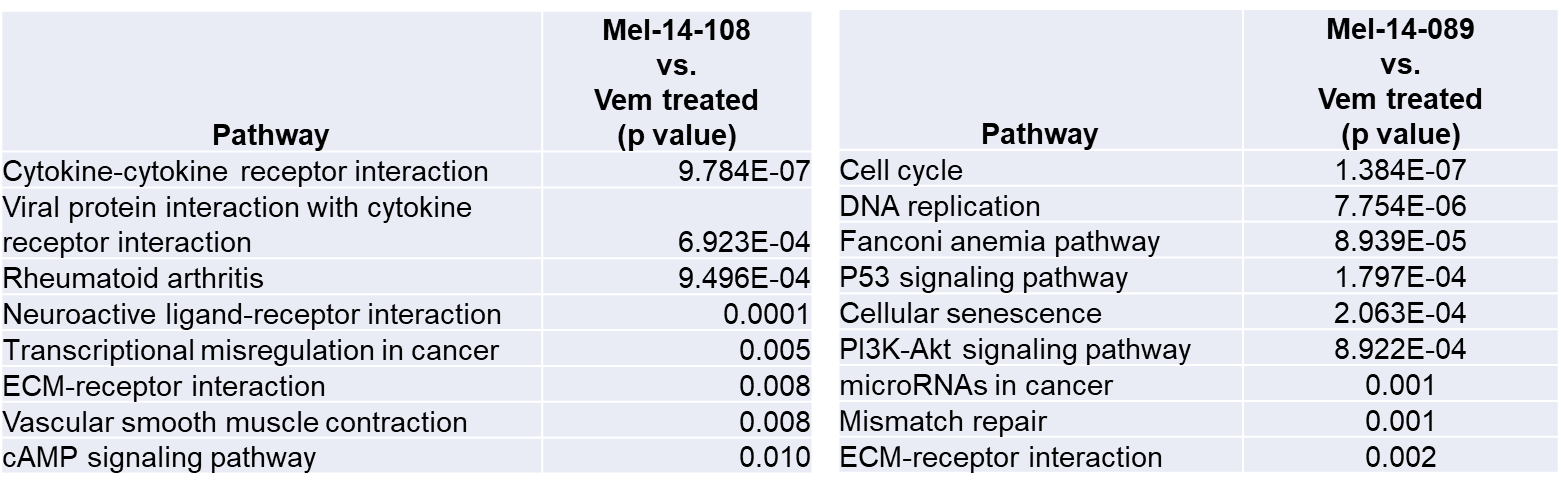
